# 5-hydroxymethylcytosine is highly dynamic across human fetal brain development

**DOI:** 10.1101/126169

**Authors:** Helen Spiers, Eilis Hannon, Leonard C. Schalkwyk, Nicholas J. Bray, Jonathan Mill

## Abstract

**Background:** Epigenetic processes play a key role in orchestrating transcriptional regulation during the development of the human central nervous system. We previously described dynamic changes in DNA methylation (5mC) occurring during human fetal brain development, but other epigenetic processes operating during this period have not been extensively explored. Of particular interest is DNA hydroxymethylation (5hmC), a modification that is enriched in the human brain and hypothesized to play an important role in neuronal function, learning and memory. In this study, we quantify 5hmC across the genome of 71 human fetal brain samples spanning 23 to 184 days post-conception.

**Results:** We identify widespread changes in 5hmC occurring during human brain development, notable sex-differences in 5hmC in the fetal brain, and interactions between 5mC and 5hmC at specific sites. Finally, we identify loci where 5hmC in the fetal brain is associated with genetic variation.

**Conclusions:** This study represents the first systematic analysis of dynamic changes in 5hmC across neurodevelopment and highlights the potential importance of this modification in the human brain. A searchable database of our fetal brain 5hmC data is available as a resource to the research community at http://epigenetics.essex.ac.uk/fetalbrain2/.

---

Human brain development is characterized by coordinated changes in gene expression mediated by a complex interaction between transcription factors[1] and epigenetic processes[2]. We recently characterized the dramatic alterations in DNA methylation (5-methylcytosine, 5mC) occurring during human neurodevelopment[3], but little is known about the role of other epigenetic modifications during this period. Of particular interest is DNA hydroxymethylation (5-hydroxymethylcytosine, 5hmC), a covalent modification of cytosine that represents an oxidised derivative of 5mC produced by the process of active DNA demethylation[4, 5]. Of note, 5hmC is present at relatively high levels in the mature central nervous system[6], and particularly enriched in the vicinity of genes with synapse-related functions[7]. Although initially hypothesized to represent an intermediate step of the DNA demethylation pathway, 5hmC is now assumed to have specific functional roles in the brain. Several specific molecular readers of 5hmC have been identified, including transcriptional regulators, chromatin modifiers and DNA damage and repair proteins[8-12]. Additionally, the ten-eleven translocation (TET) family of proteins that catalyze the conversion of 5mC into 5hmC have been implicated in neuronal differentiation and function[13]; studies manipulating TET activity suggest that 5hmC plays a role in learning and memory, hippocampal neurogenesis, and neuronal activity-regulated gene expression[14-16]. Recently, 5hmC has been found to mark regulatory regions of the genome in the murine and human fetal brain[2, 17] and increase rapidly in abundance postnatally, concurrent with neuronal maturation and synaptogenesis[18-20]. Notably, there is growing evidence to suggest that abnormal regulation of 5hmC contributes to the etiology of several neurodevelopmental disorders including autism spectrum disorders[21] and schizophrenia[22].

Importantly, many of the standard methods used to quantify 5mC, including those based on sodium bisulfite (BS) conversion of DNA, are unable to discriminate between 5mC and 5hmC[10, 23], with important implications for the interpretation of published DNA methylation datasets from brain. Oxidative bisulfite (oxBS) conversion of genomic DNA, which involves the oxidation of 5hmC to 5-formylcytosine (5fC) prior to standard BS conversion, enables the accurate quantification of 5mC, allowing 5hmC to be measured by proxy at base-pair resolution[24-26]. In this study, we use this approach in combination with the Illumina 450K HumanMethylation array (“450K array”) to undertake the first systematic study of 5hmC in the developing human brain, profiling 71 fetal samples ranging from 23 to 184 days post-conception (DPC). We identify widespread changes in 5hmC during human brain development, with differentially hydroxymethylated positions (DHPs) and regions (DHRs) becoming both hyper- and hypo-hydroxymethylated with fetal age. As a resource to the scientific community a searchable database of our fetal brain 5hmC data is available at http://epigenetics.essex.ac.uk/fetalbrain2/.

## RESULTS

### Autosomal distribution of 5hmC in the developing human brain

We quantified 5hmC across the genome by subtracting oxBS-generated 450K array profiles from those generated following BS-conversion performed in parallel (Δβ_BS-oxBS_). We set a stringent threshold for calling 5hmC (Δβ_BS-oxBS_ > 0.036) based on the 95th percentile of negative Δβ_BS-oxBS_ values across all profiled samples, with sites below this threshold characterized as having “undetectable” 5hmC (see **Methods** section). Consistent with previous observations in young neurons[18, 19, 27], we observed overall low levels of global 5hmC in the fetal brain (mean level of 5hmC across all 411,325 probes passing our stringent quality-control metrics = 2.97% (SD = 5.16%)). Approximately a quarter (n = 103,063 (25.64%)) of all autosomal probes included in the analysis were characterized by non-detectable 5hmC (i.e. Δβ_BS-oxBS_ < 0.036) in all 71 fetal brain samples examined (Fig.1a and **Supplementary Table 1**); these sites were significantly enriched in CpG islands and other promoter regulatory regions including the transcription start-site (TSS), 5’UTR and 1^st^ Exon (**Supplementary Table 2**). In contrast, a small number of autosomal sites (n = 5,061 (1.26%)) were characterized by detectable 5hmC in all 71 samples; these sites were significantly enriched in CpG island shores and shelves, regions flanking the TSS and gene bodies (**Supplementary Table 2**). Notably, the majority of probes were characterized by detectable 5hmC in only a subset of samples (Fig. 1a), indicating that the presence and distribution of 5hmC is highly variable during fetal brain development.

**Figure 1:**
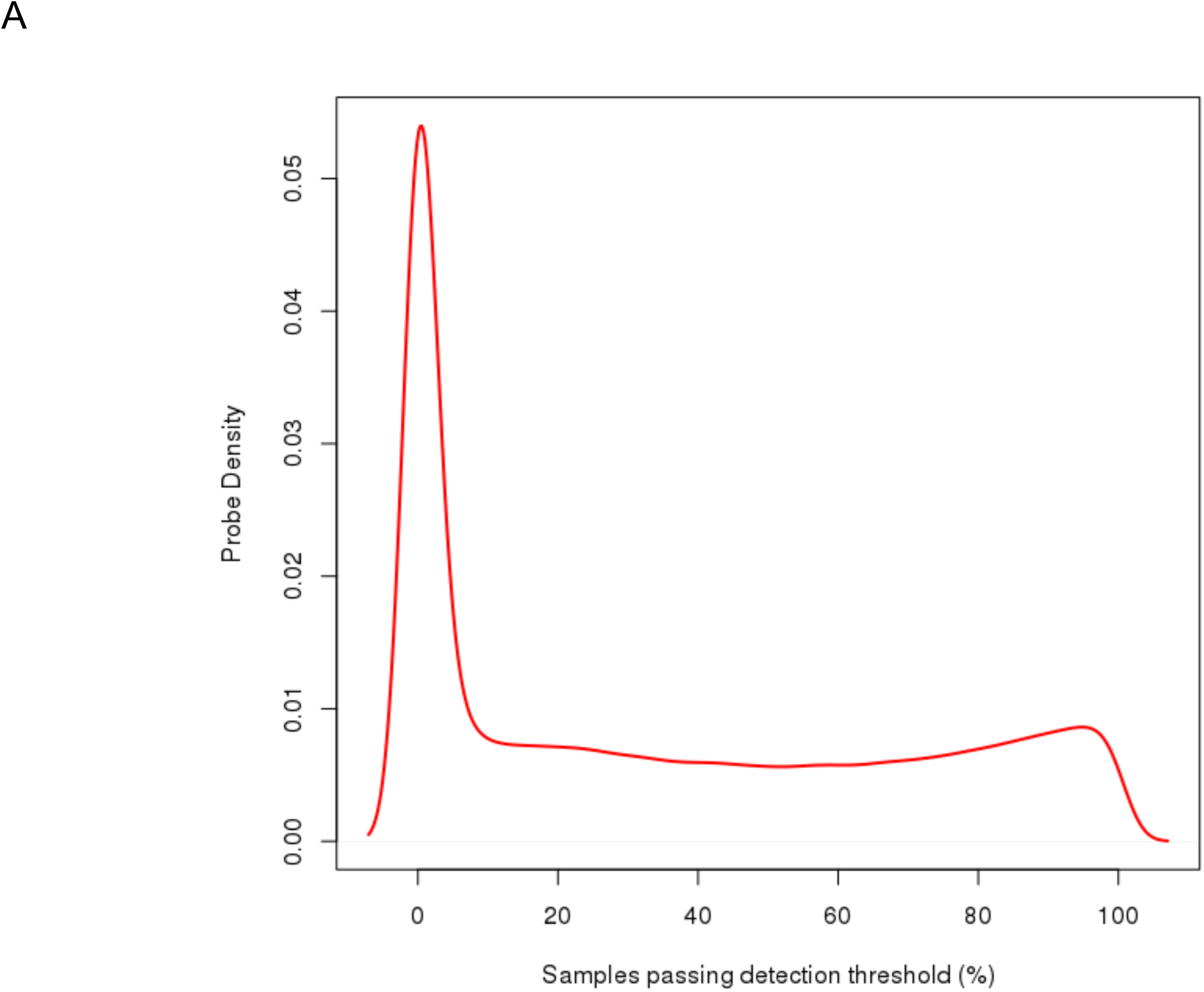
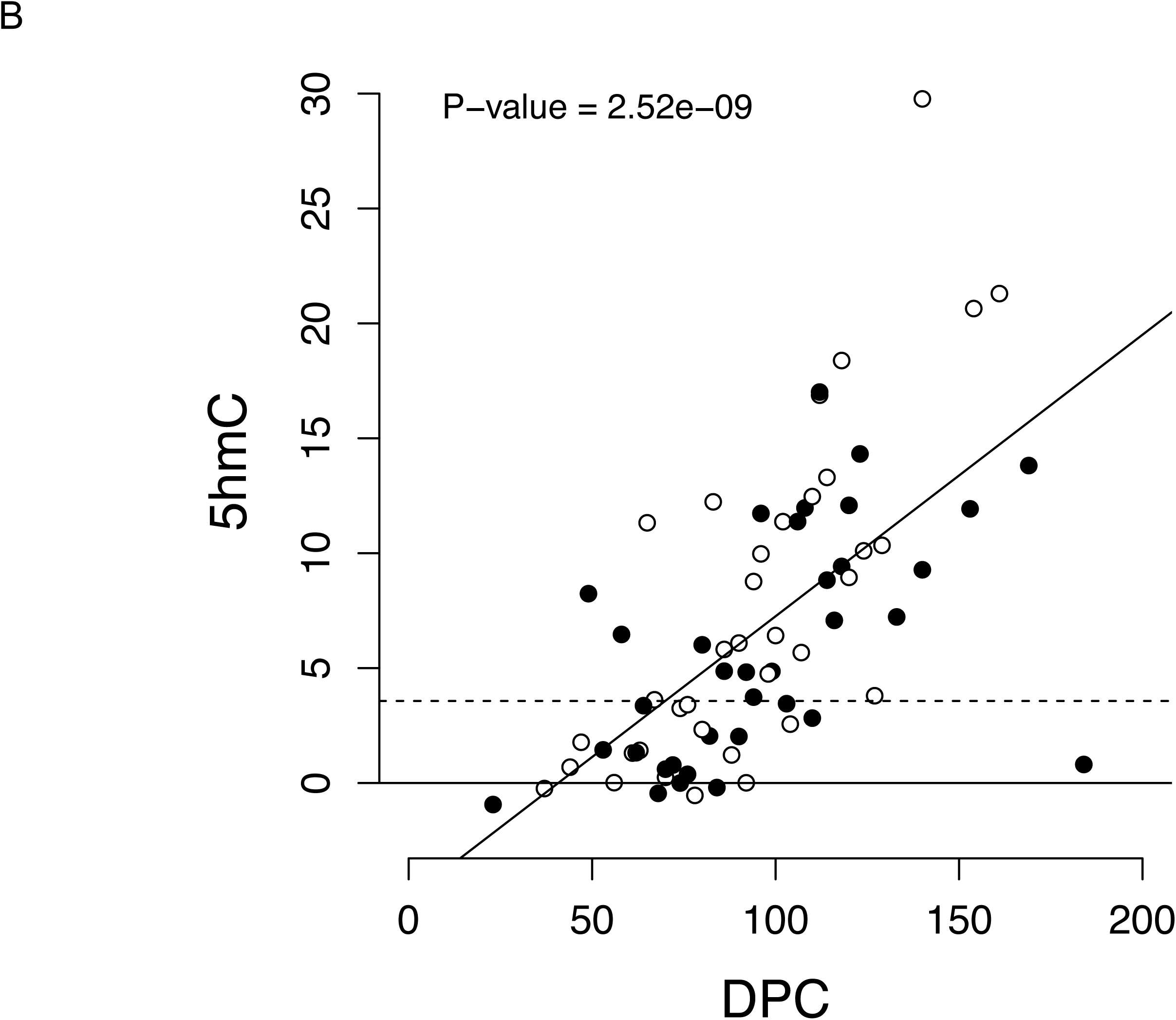
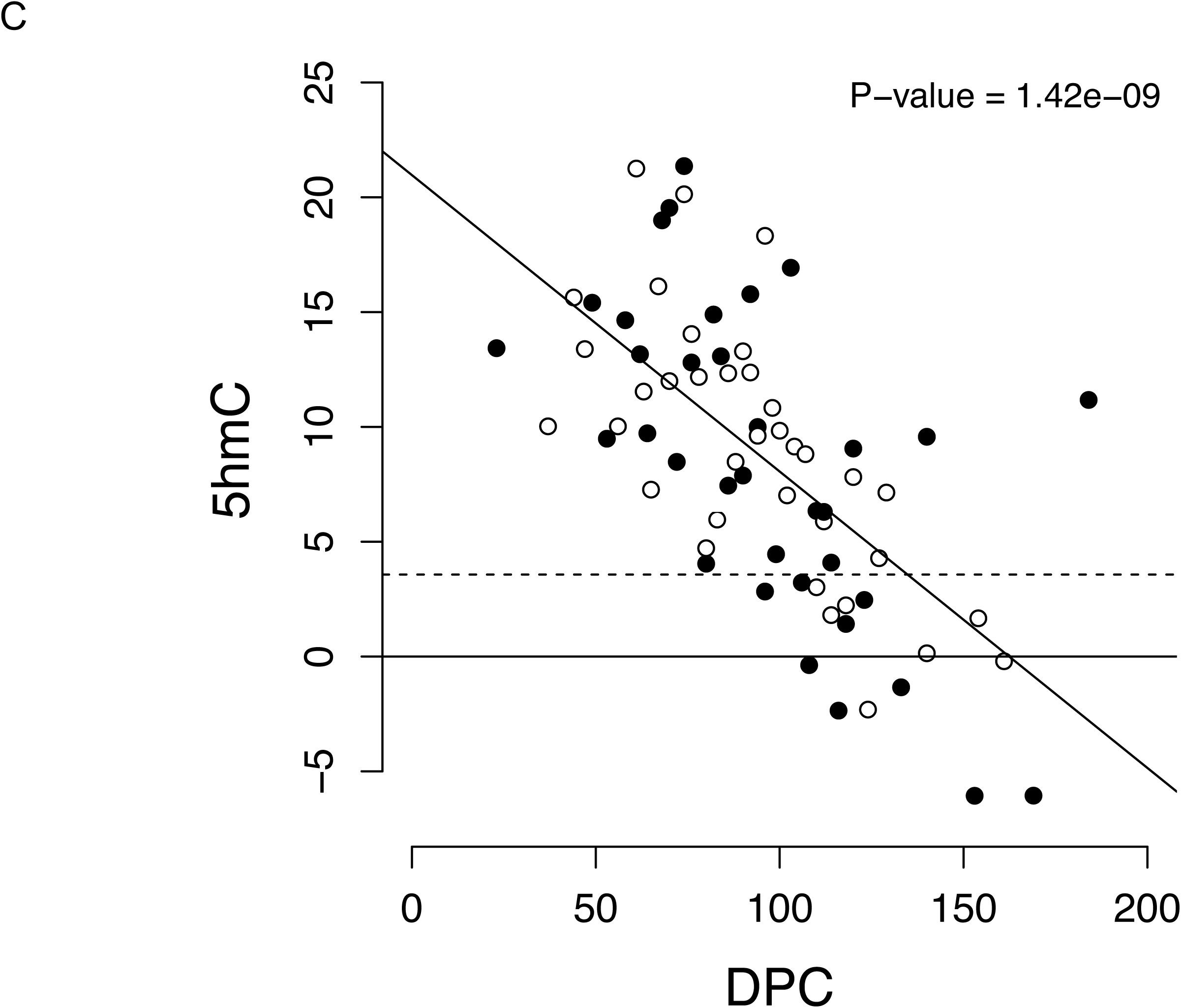
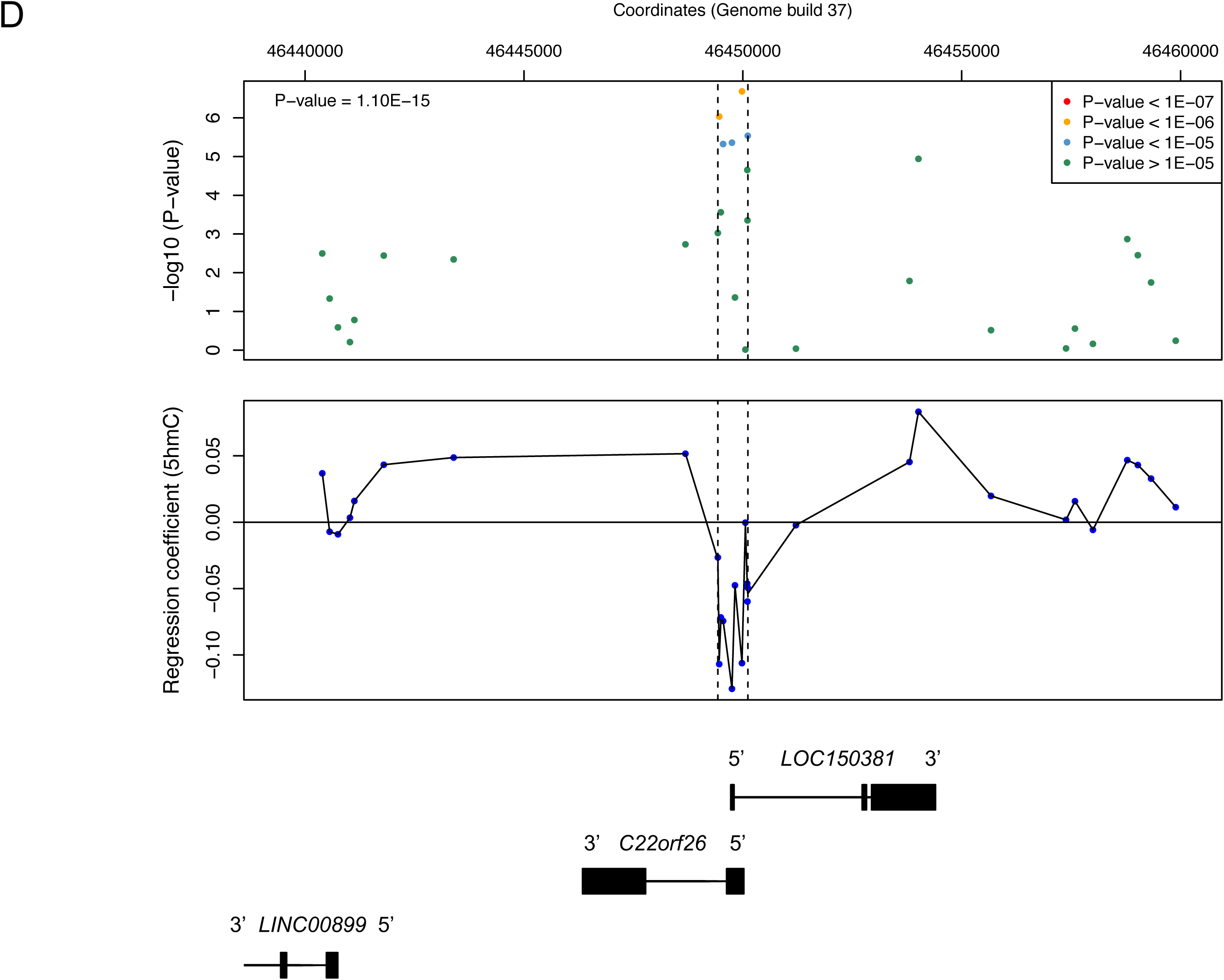
DNA hydroxymethylation is highly dynamic across human fetal brain development. (A) The presence or absence of detectable 5hmC at individual sites is highly variable amongst the 71 fetal brain samples profiled in this study. Shown is the proportion of probes with detectable 5hmC as a function of the fetal brain samples profiled. Only a small number of probes are characterized by ubiquitously detectable 5hmC in all samples, highlighting considerable inter-individual heterogeneity at individual probes. (B) The top-ranked dDHP characterized by increasing 5hmC with days post-conception (DPC) is cg25069807, annotated to *TK1* on chromosome 17 (*P* = 2.52E-09). (C) The top-ranked dDHP characterized by decreasing 5hmC with brain development is cg22960621, annotated to *NLGN1* on chromosome 3 (*P* = 1.42E-09). Males are depicted in black. The dashed line indicates values above which 5hmC is considered to be reliably detected (Δβ_BS-oxBS_ > 0.036). The full list of 2,181 dDHPs (*P* < 5.00E-05) is presented in **Supplementary Table 5**. (D) The top-ranked dDHR associated with fetal brain development spans a 685bp region (Chr22: 46449430 –46450115) encompassing the first exon of *C22orf26* and *LOC150381*. The dDHR is denoted by dashed vertical lines. Chromosomal coordinates correspond to human genome build GRCh37/hg19. See **Supplementary Table 8** for a full list of dDHRs associated with human brain development.

Across the 298,972 autosomal probes (74.36%) characterized by detectable 5hmC (i.e. Δβ_BS-oxBS_ > 0.036) in at least one fetal brain sample, the mean level of 5hmC was 4.16% (SD 5.43%) across all fetal brain samples (**Supplementary Table 3**). In agreement with previous studies[26, 28-31], we observed an inverse relationship between CpG density and mean levels of 5hmC at these sites, with significantly lower levels in CpG islands (1.56%, *P* < 1.00E-200) and higher levels in sites not located within CpG islands, shores or shelves (5.20%, *P* < 1.00E-200) compared to the average across all sites characterized by detectable 5hmC in at least one fetal brain sample (**Supplementary Fig. 1**; **Supplementary Table 3**). 5hmC was significantly enriched at 450K array probes in gene bodies (4.37%, *P* < 1.00E-200) and 3’UTRs (4.74%, *P* < 1.00E-200), and significantly depleted at 450K array probes located within 1500 bp (3.93%, *P* < 1.00E-200) and 200 bp (2.14%, *P* < 1.00E-200) upstream of the transcription start site (TSS), 5’UTRs (3.85%, *P* < 1.00E-200), and 1^st^ exons (2.24%, *P* < 1.00E-200) (**Supplementary Fig. 1**; **Supplementary Table 3**), concurring with previous observations[19, 26, 28-30].

### Human fetal brain development is associated with dynamic changes in 5hmC at specific loci

Our previous study quantified BS-converted DNA to identify changes in total DNA modifications (i.e. 5mC *and* 5hmC) across fetal brain development in an overlapping set of fetal brain samples[3], and was therefore not able to distinguish between 5mC and 5hmC. For the developmentally differentially modified positions (dDMPs) identified in our original study that were included in our final dataset (n = 28,330 probes), we found a highly-significant correlation of effect sizes across both BS datasets (r = 0.99; *P*-value < 1.00E-200) (**Supplementary Fig. 2** and **Supplementary Fig. 3**), confirming the robust nature of our previous DNA modification estimates. We explored the extent to which changes at these sites, previously attributed to changes in 5mC, were confounded by levels of 5hmC. There was a strong correlation (r = 0.97, *P*-value < 1.00E-200) between the BS (5mC + 5hmC) and oxBS (5mC) effect sizes (**Supplementary Fig. 4**) at these sites, indicating that the DNA modification dynamics previously described[3] are likely to primarily reflect changes in 5mC. The majority of these sites (n = 28,255, 99.74%), however, were characterized by detectable 5hmC in at least one sample and there was a significant positive correlation (r = 0.36; *P*-value < 1.00E-200) (**Supplementary Fig. 5**) between the BS (5mC + 5hmC) and BS-oxBS (5hmC) effect sizes, indicating a contribution of dynamic 5hmC across brain development at these sites.

We next used a linear model to detect sites where levels of 5hmC changed across brain development (see Methods), identifying 62 developmentally differentially hydroxymethylated positions (dDHPs) at a stringent Bonferroni-corrected significance threshold (*P*-value < 1.67E-07) (**Supplementary Table 4**), representing 0.02% of the 298,972 autosomal probes tested. At a more relaxed “discovery” significance threshold (*P*-value < 5E-05), we identified 2,181 sites (0.73%) characterized by changes in 5hmC during brain development (**Supplementary Table 5**). The top-ranked sites characterized by increasing 5hmC across brain development (henceforth referred to as hyperhydroxymethylated dDHPs) and decreasing 5hmC across brain development (henceforth referred to as hypohydroxymethylated dDHPs) are shown in Fig. 1b and Fig. 1c, respectively. Among the “discovery” dDHPs we observed a significant enrichment of hypohydroxymethylated autosomal dDHPs (n = 1,278 (58.60%)) compared to hyperhydroxymethylated dDHPs (n = 903 (41.40%)) (enrichment hyperhydroxymethylated OR = 0.71, *P*-value = 1.29E-08) (**Supplementary Table 6**). We next used comb-P[32] to identify spatially correlated regions of developmentally dynamic 5hmC (or developmentally differentially hydroxymethylated regions (dDHRs)). A total of 254 dDHRs (Šidák-corrected *P-*value < 0.05) were identified, spanning an average of 5.43 probes and 551.30 base pairs (SD = 362.97) (**Supplementary Table 7**). The top-ranked dDHR, encompassing 11 CpG sites spanning the first exon of *C22orf26* and *LOC150381* on chromosome 22, is characterized by a significant reduction in 5hmC across brain development (Šidák-corrected *P*-value = 1.10E-15) (Fig. 1d). A full list of dDHRs is presented in **Supplementary Table 8**.

### Sites characterized by changes in 5hmC across brain development are not distributed equally across genomic features

Although the frequency of dDHPs is relatively consistent across autosomes, chromosome 19 is notably depleted of dDHPs (relative enrichment = 0.62, *P*-value = 3.08E-05) (**Supplementary Table 9**). The distribution of dDHPs, however, is not equal across annotated genic features (**Supplementary Table 6**, **Supplementary Table 10**), being significantly depleted in promoter regulatory regions, including CpG islands (percentage of significant probes = 0.32%, relative enrichment = 0.43, *P*-value = 1.43E-33), and enriched at sites not annotated to CG-rich features (CpG islands, shores and shelves) (percentage of significant probes = 0.92%, relative enrichment = 1.26, *P*-value = 2.27E-10) (**Supplementary Fig. 6** and **Supplementary Fig. 7**). There is also a significant enrichment of dDHPs in annotated DNase I hypersensitivity sites (percentage of significant probes = 0.96%, relative enrichment 1.32, *P*–value = 2.62E-13) (**Supplementary Table 6**) and probes associated with transcription factor binding sites (TFBSs)[33] (percentage of significant probes = 0.79%, relative enrichment = 1.09, *P*-value = 4.41E-02), with specific TFBS motifs being specifically enriched for dDHPs (see **Supplementary Table 11**). Sites associated with specific alternative transcription event domains were significantly depleted for dDHPs (**Supplementary Fig. 8**; **Supplementary Table 12**) including alternative 3’ splice sites (percentage of significant probes = 0.34%, relative enrichment = 0.46, *P*-value = 1.51E-02) and constitutive exons (percentage of significant probes = 0.44%, relative enrichment = 0.59, *P*-value = 4.70E-06).

### There is a complex relationship between 5mC and 5hmC across human brain development

It is not yet definitively known whether 5hmC represents a stable or transient DNA modification[34] and its relationship to 5mC has not been explored in a large number of samples. Our data show that levels of 5hmC in the fetal brain are highest at loci characterized by intermediate levels of 5mC (**Supplementary Fig. 9**), concurring with the results of previous analyses[26, 29]. To further delineate the relationship between 5mC and 5hmC during brain development we compared effect sizes (regression coefficients reflecting change in DNA modification with development) for all dDHPs passing our ‘discovery’ significance threshold (n = 2,181) (Fig. 2a). dDHPs could be split into three broad groups; one characterized by changes in 5mC paralleling those seen for 5hmC (e.g. Fig. 2b-c), a second where changes occurred in the opposite directions across the two DNA modifications (e.g. Fig. 2d-e), and a third characterized by changes in 5hmC, but no changes in 5mC (e.g. Fig. 2f). Of note, nearly half (n = 1,059; 48.56%) of the dDHPs tested where characterized by a significant interaction (*P*-value < 2.29E-05) between trajectories of 5mC and 5hmC across brain development.

**Figure 2:**
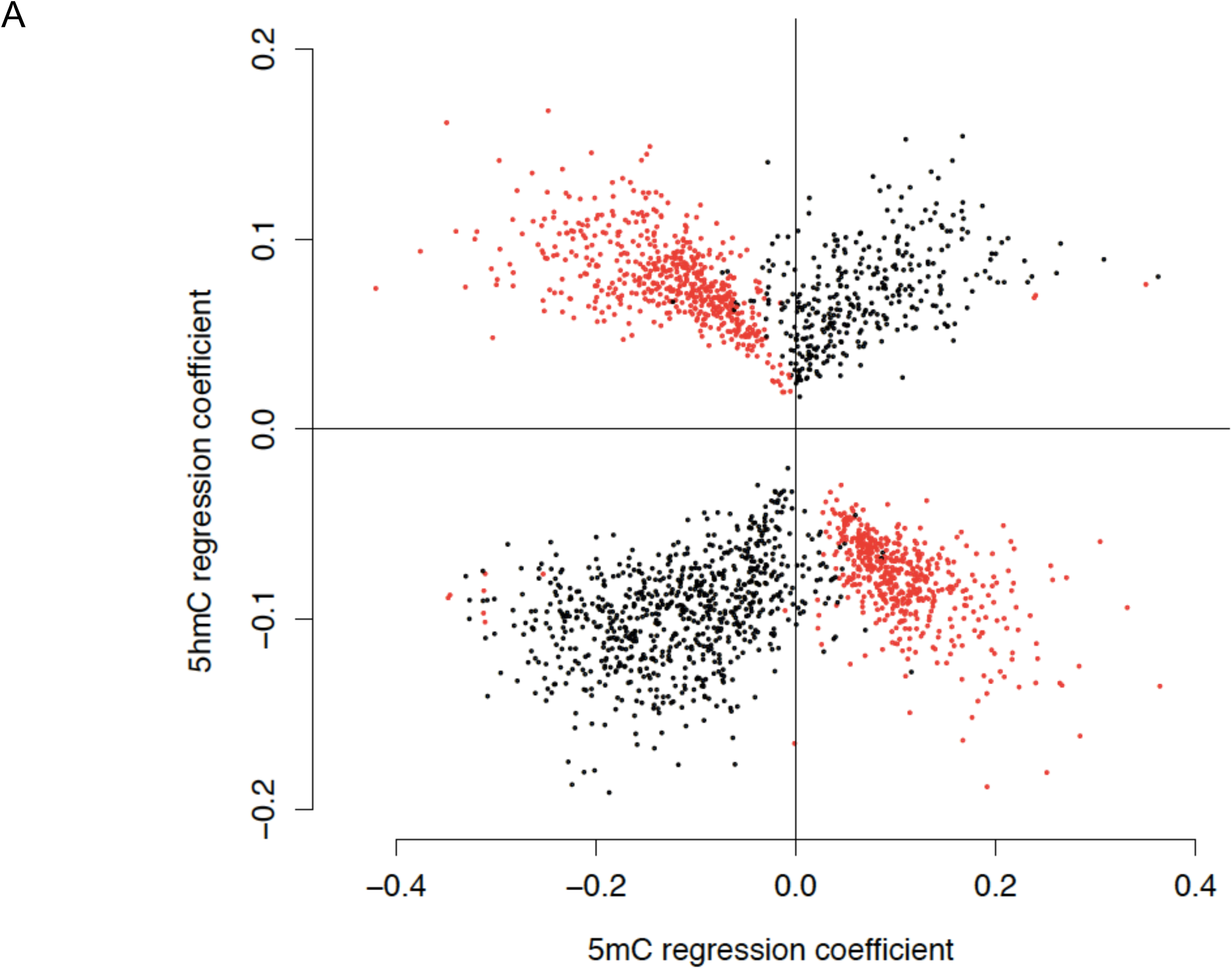
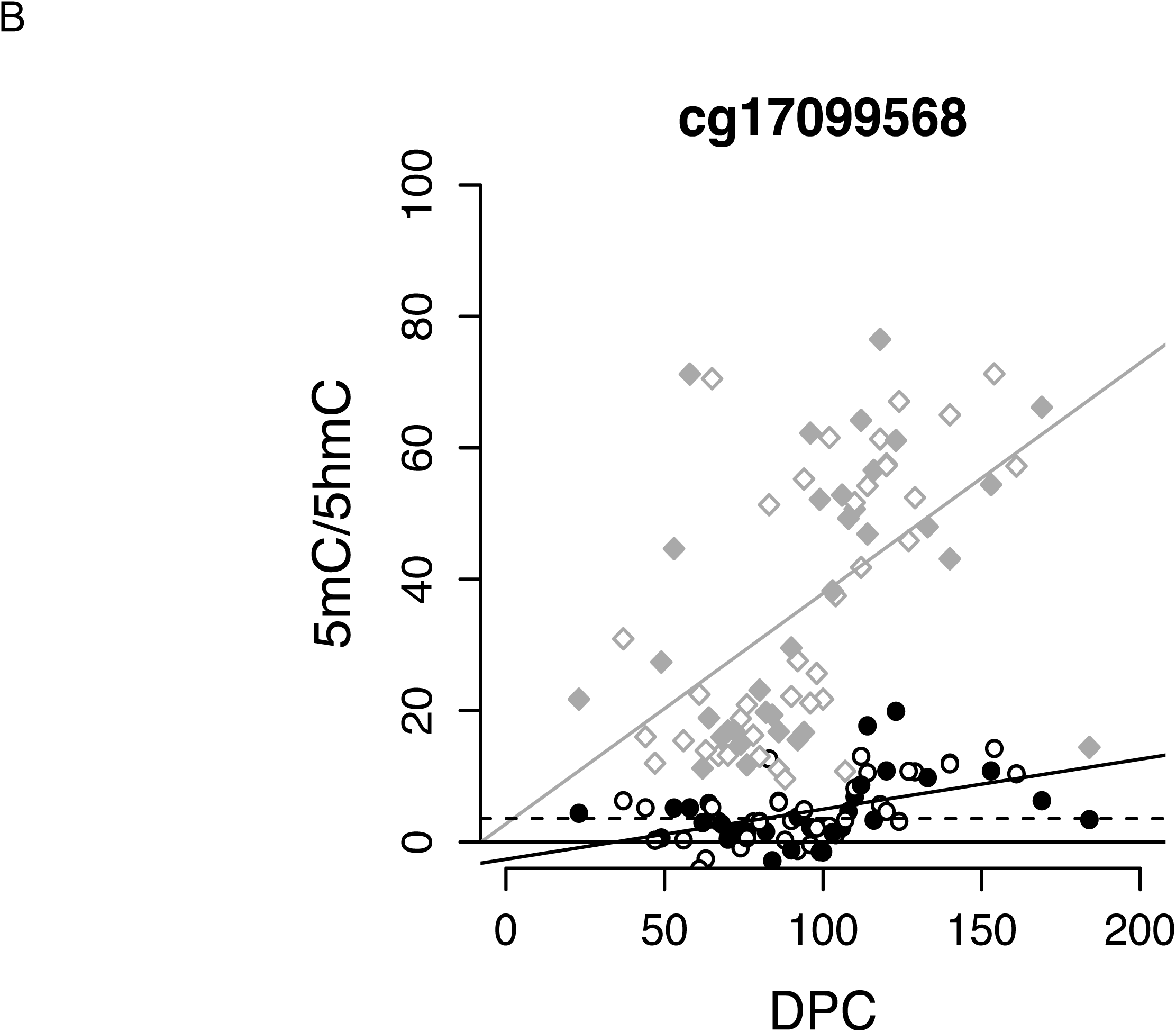
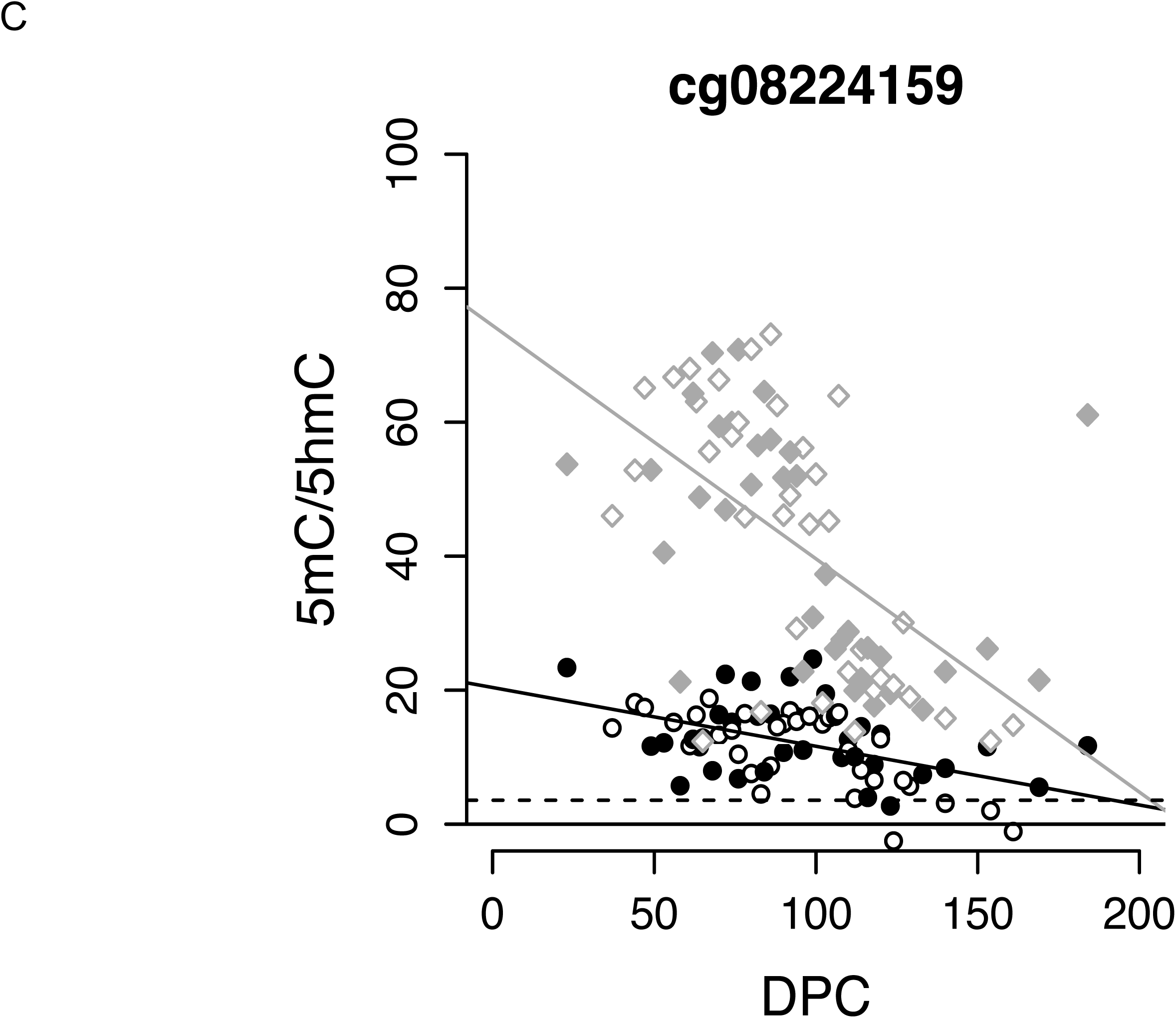
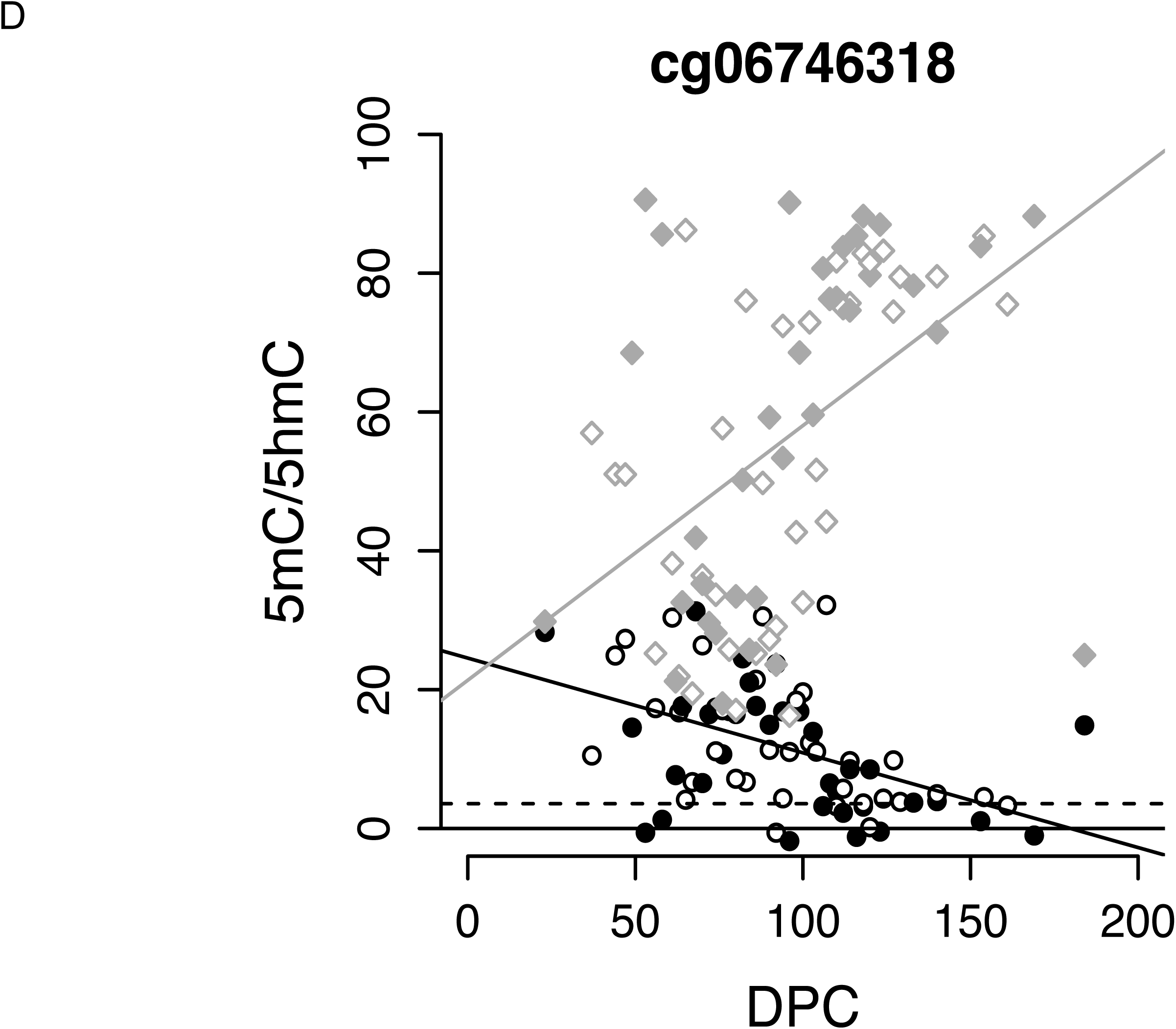
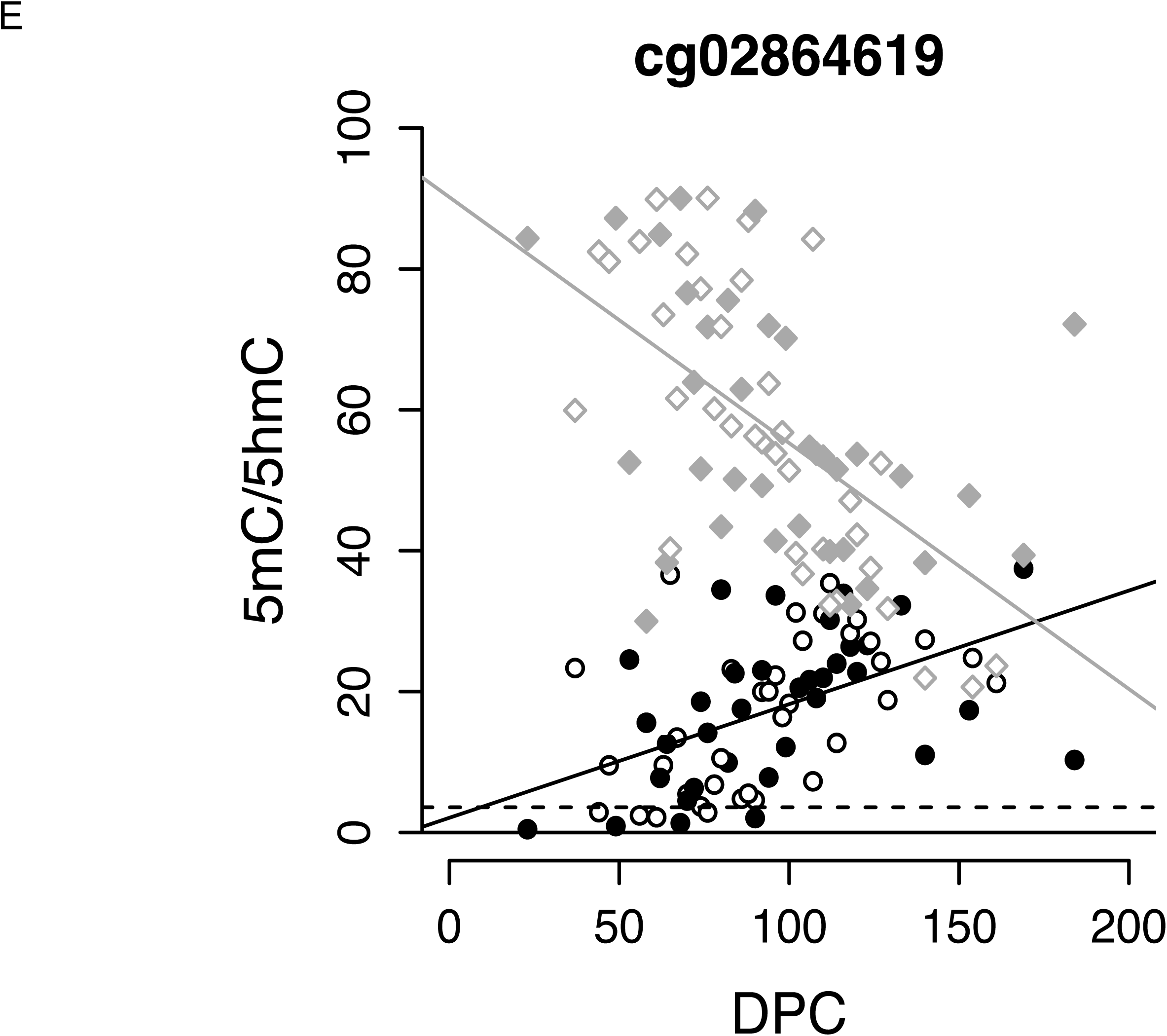
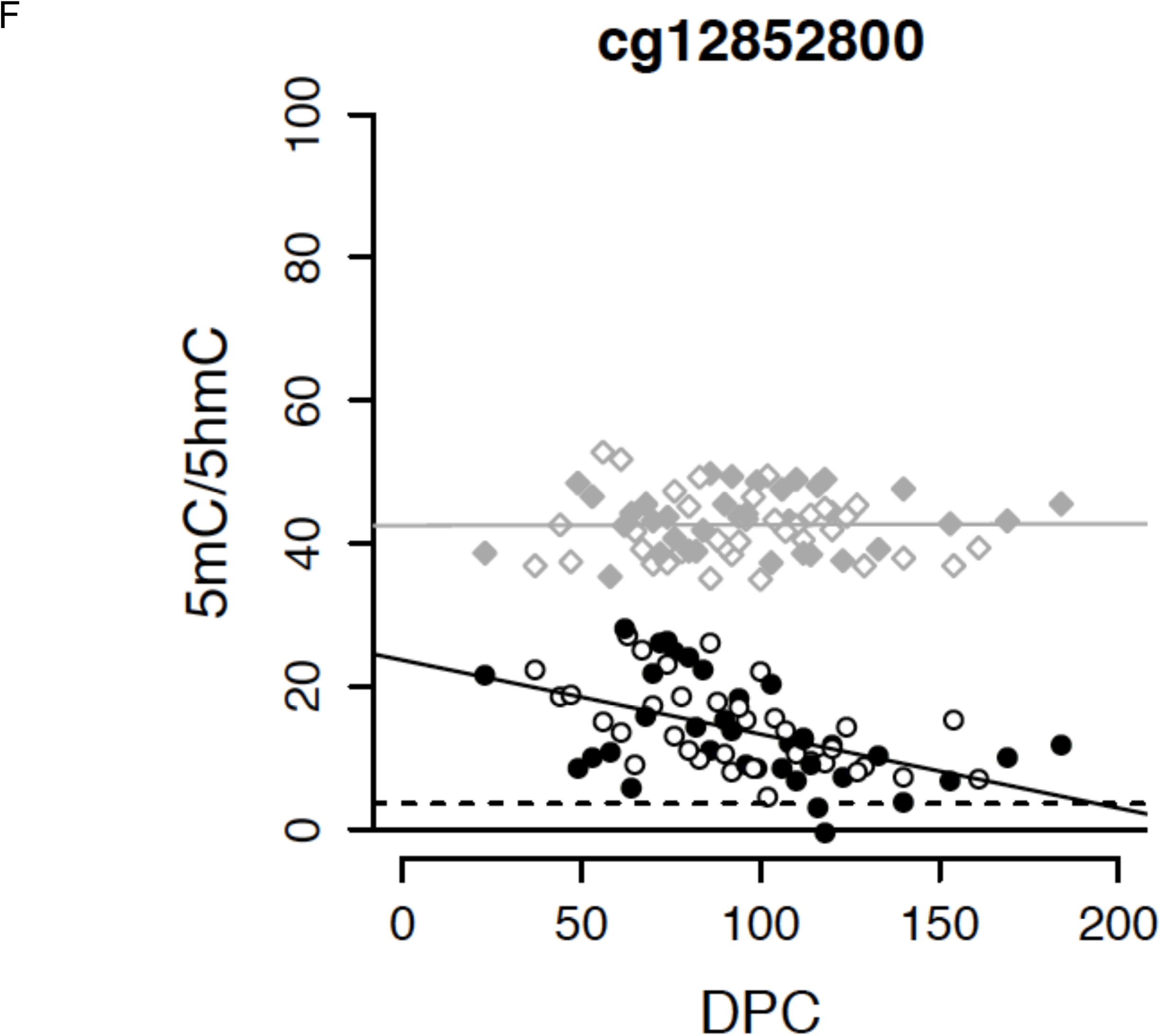
There is a complex interaction between 5hmC and 5mC across human brain development. (A) Scatterplot showing the regression coefficients for 5hmC and 5mC for each of the 2,181 dDHPs (*P* < 5.00E-05). Nearly half of the dDHPs (n = 1059, 48.56%) are characterized by a significant interaction (*P* < 2.29E-05) between 5hmC and 5mC across brain development (significant sites are shown in red). Examples of sites at which fetal brain development is associated with (B) increases in both 5hmC and 5mC, (C) decreases in both 5hmC and 5mC, (D) an increase in 5mC but a decrease in 5hmC, (E) a decrease in 5mC but an increase in 5hmC and (F) a change in 5hmC but no alteration in 5mC. 5mC data shown as diamonds, 5hmC data shown as circles. Male samples are indicated by a filled diamond or circle.

### Sex differences in the human fetal brain hydroxymethylome

Although the majority of autosomal chromosomes are characterized by a similar mean level of 5hmC (**Supplementary Table 13**) (mean 5hmC across 298,972 autosomal probes = 4.16%), levels are significantly lower on the X-chromosome (mean 5hmC = 2.00%, *P* < 1.00E-200), especially in females (mean 5hmC across X-chromosome in females = 1.40%, *P* < 1.00E-200). A total of 427 probes (0.14% of the 307,810 autosomal and X-linked probes assessed) displayed significantly different levels of 5hmC between male and females (Bonferroni-significance level of *P*-value < 1.62E-07) (Fig. 3a). All were located on the X-chromosome, suggesting that, like 5mC, 5hmC is likely to play a role in X-chromosome dosage compensation mechanisms. In contrast to 5mC, however, which is elevated on the inactive female X-chromosome, the majority of these sites (n = 404; 94.61%) were characterized by a higher level of 5hmC in males (Fig. 3b, Fig. 3c and **Supplementary Fig. 10**). Higher 5hmC in males versus females is observed across canonical gene features on the X chromosome, however this is not seen on the autosomes (Fig. 3d). A total of 144 fetal brain DHRs were identified between males and females using *comb-P* (Šidák-corrected P-value < 0.05, number of probes ≥ 3) (**Supplementary Table 14**). All were located on the X-chromosome and the top-ranked sex-associated DHR is shown in **Supplementary Fig. 11**.

**Figure 3:**
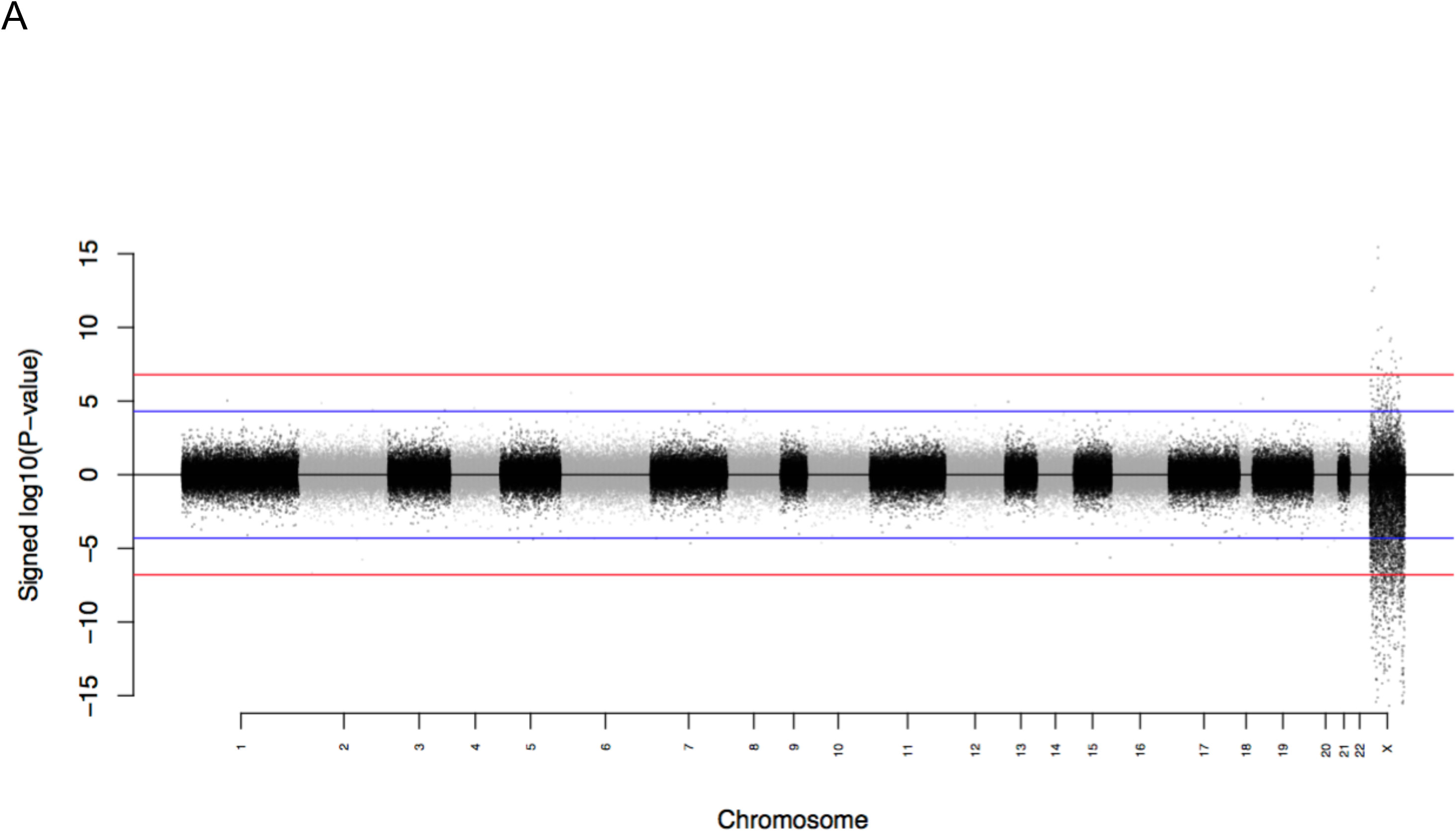
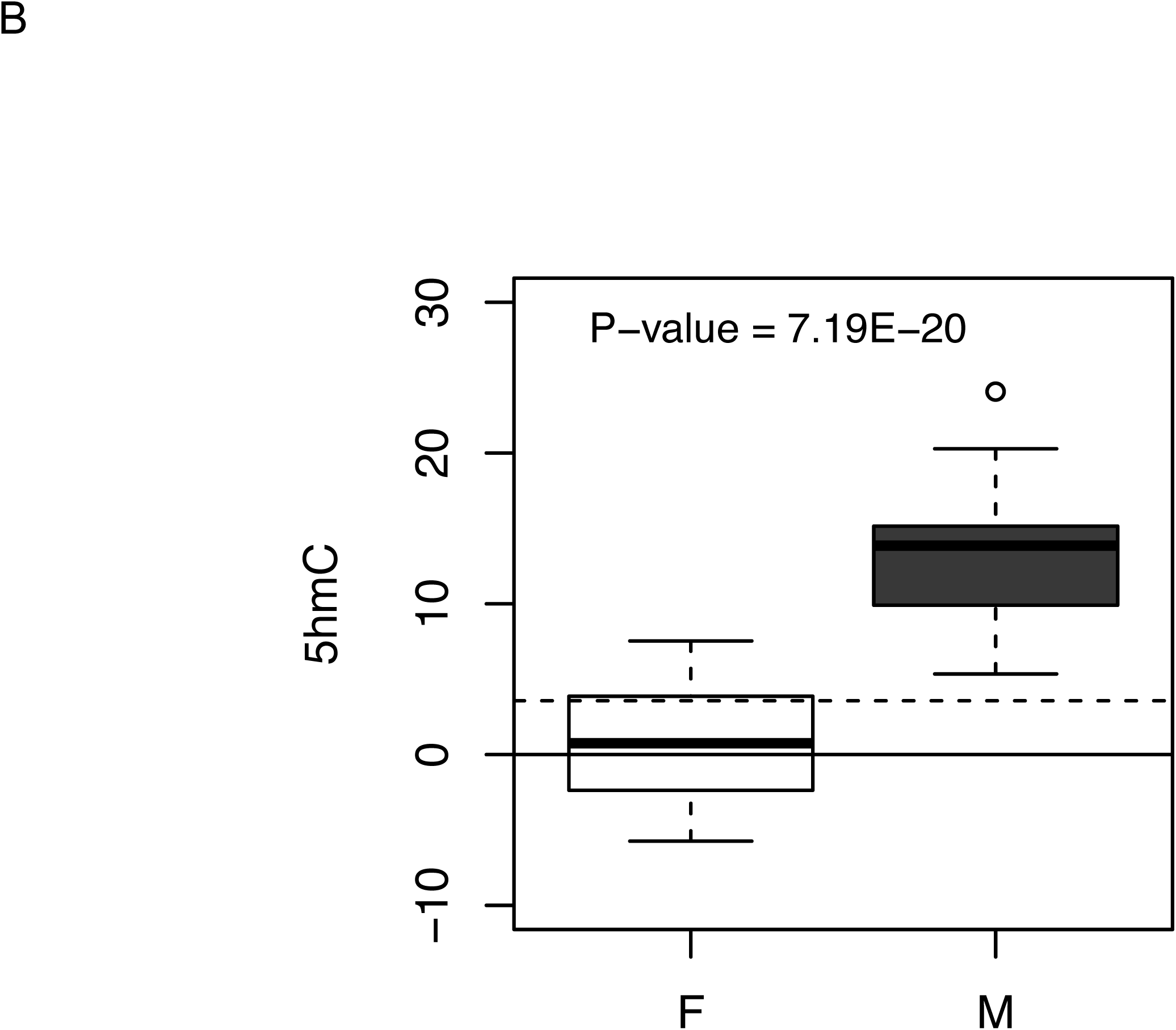
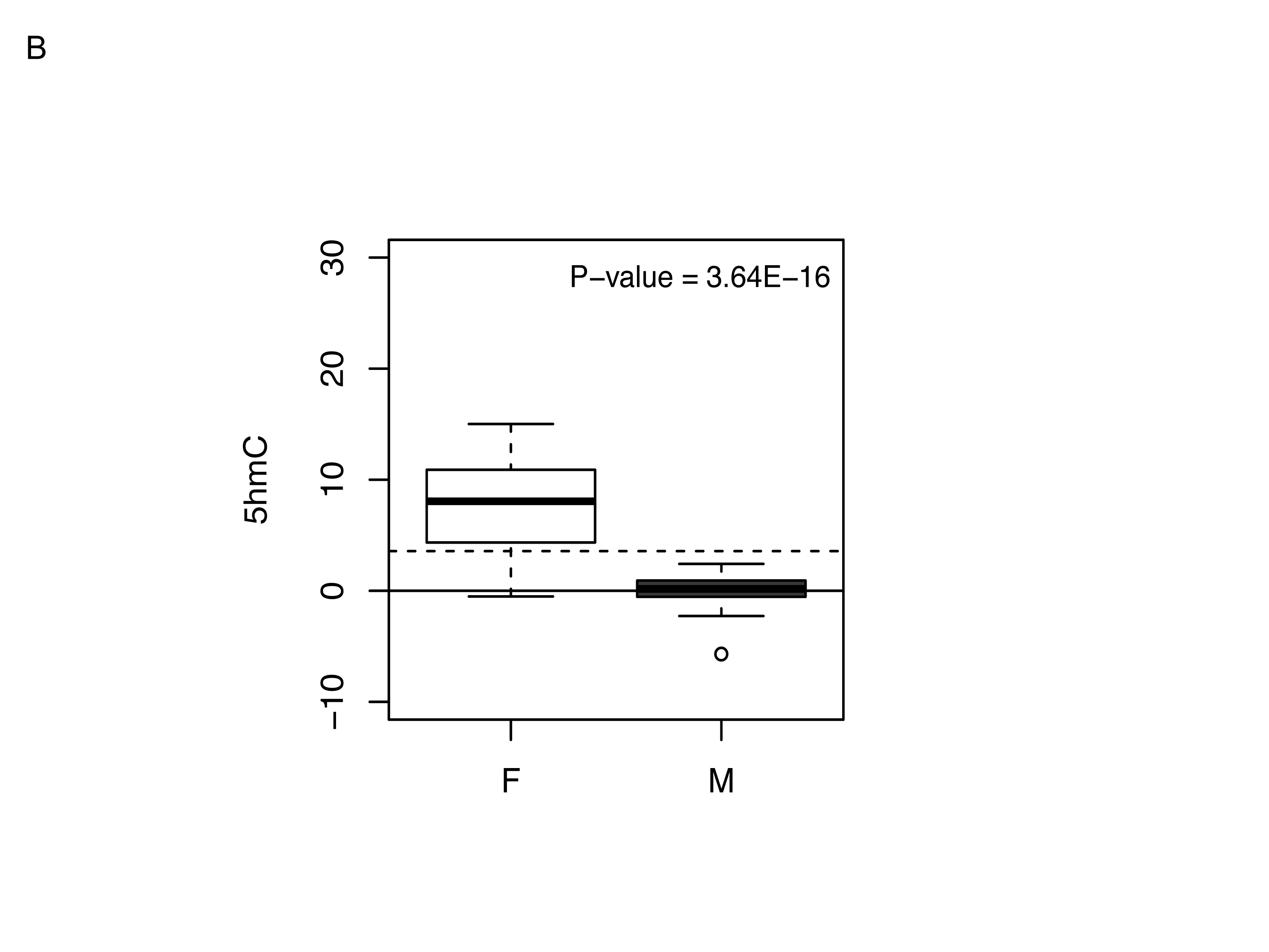
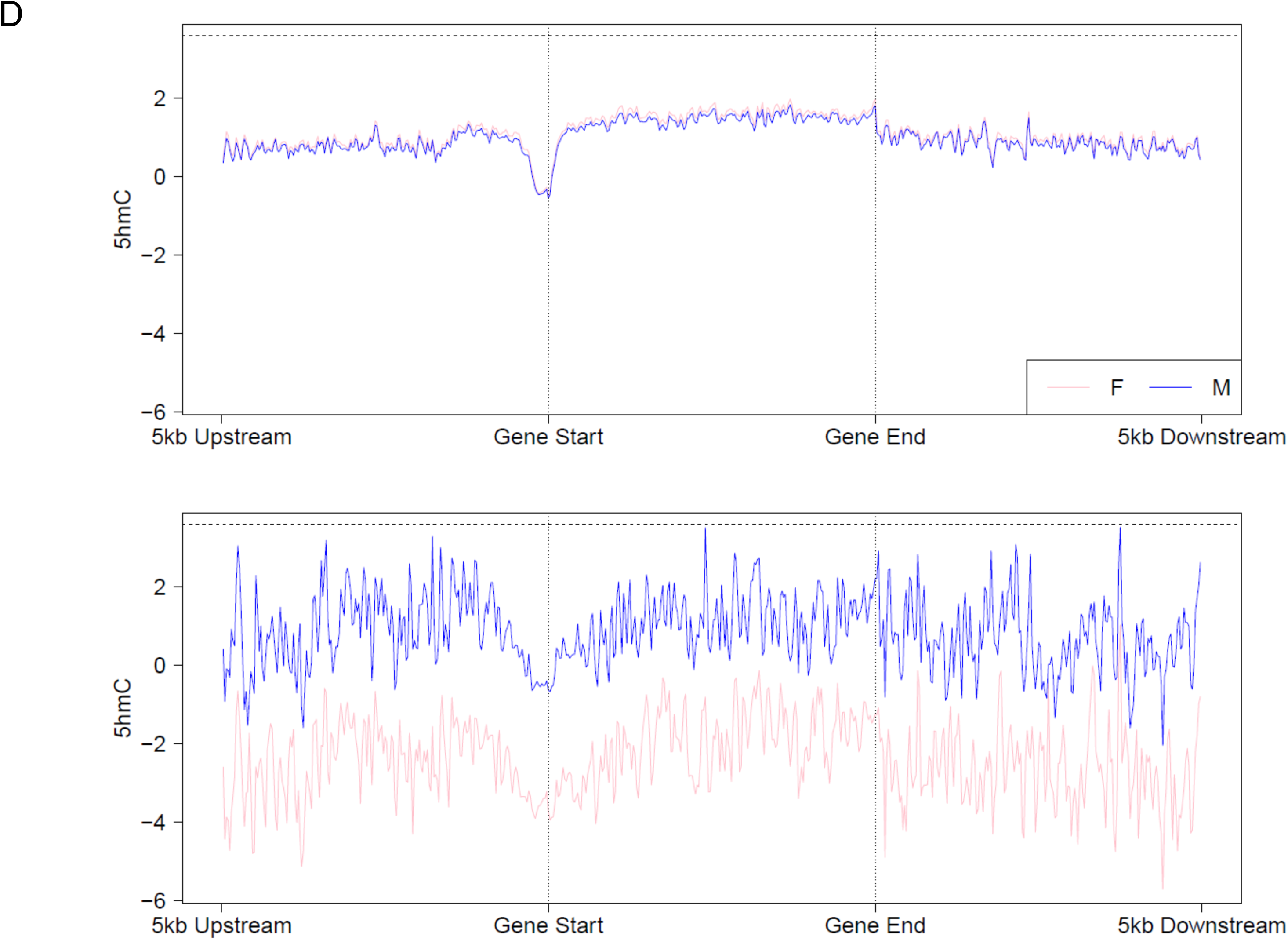
Levels of 5hmC in the human fetal brain are significantly lower on the X-chromosome in females than males. (A) Manhattan plot showing the distribution of sites characterized by different levels of 5hmC between males and females. *P*-value corresponds to association with sex. Points above the X-axis indicate higher 5hmC in females, with those below the X-axis indicating higher 5hmC in males. The red line represents a Bonferroni-corrected significance threshold (*P* < 1.62E-07) for the 307,810 autosomal and X chromosome probes analyzed. The blue line corresponds to our “discovery” *P*-value threshold (P < 5E-05). (B) The top-ranked sex-associated DHP characterized by greater 5hmC in males is cg07806797, annotated to ZNF185 on the X-chromosome (P = 7.19E-20). (C) The top-ranked sDHP characterized by greater 5hmC in females is cg2678647, annotated to USP11 on the X-chromosome (P = 3.64E-16). (D) Mean 5hmC across canonical gene features is shown for males and females (pink and blue, respectively) for autosomes (top panel) and the X-chromosome (bottom panel).

### Modules of co-hydroxymethylated loci are identified in the developing human brain

We next employed weighted gene co-hydroxymethylation network analysis (WGCNA)[35] to undertake a systems level characterization of the 5hmC changes associated with human brain development. WGCNA identified a total of 32 discrete modules of co-hydroxymethylated sites (**Supplementary Table 15**). The first principal component of each module, the “module eigengene”, was used to assess the module relationship with fetal brain development (Fig. 4a). Seven modules were significantly correlated with brain development (*P*-value < 1.56E-03) (**Supplementary Table 15**; Fig. 4b; **Supplementary Fig. 12**). Module membership within these modules was highly correlated with probe significance (**Supplementary Fig. 13**), indicating a clear relationship between 5hmC at the core members of each module and fetal brain development (Fig. 4c). To assess whether these modules are biologically meaningful, functional enrichment analyses were performed on the top 1,000 probes from the seven DPC-associated modules (**Supplementary Table 16**). Highly significant gene ontology (GO) terms were identified for the DPC associated modules including “dendrite morphogenesis” (“blue” module; *P*-value = 3.22E-11), “negative regulation of synaptic transmission” (“black” module; *P*-value = 6.29E-09), “negative regulation of axon extension involved in axon guidance” (“red” module; *P*-value = 3.64E-09) and “embryonic organ morphogenesis” (“green” module, *P*-value = 1.79E-11), suggesting that the large changes in 5hmC across fetal brain development contribute to the regulation of relevant biological processes. Additionally, the sex-associated “lightcyan” module was significantly enriched for terms relevant to brain development including “dendritic spine development” (*P*-value = 8.86E-05), suggesting that sex differences in 5hmC may contribute to sex-specific neurodevelopmental processes.

**Figure 4:**
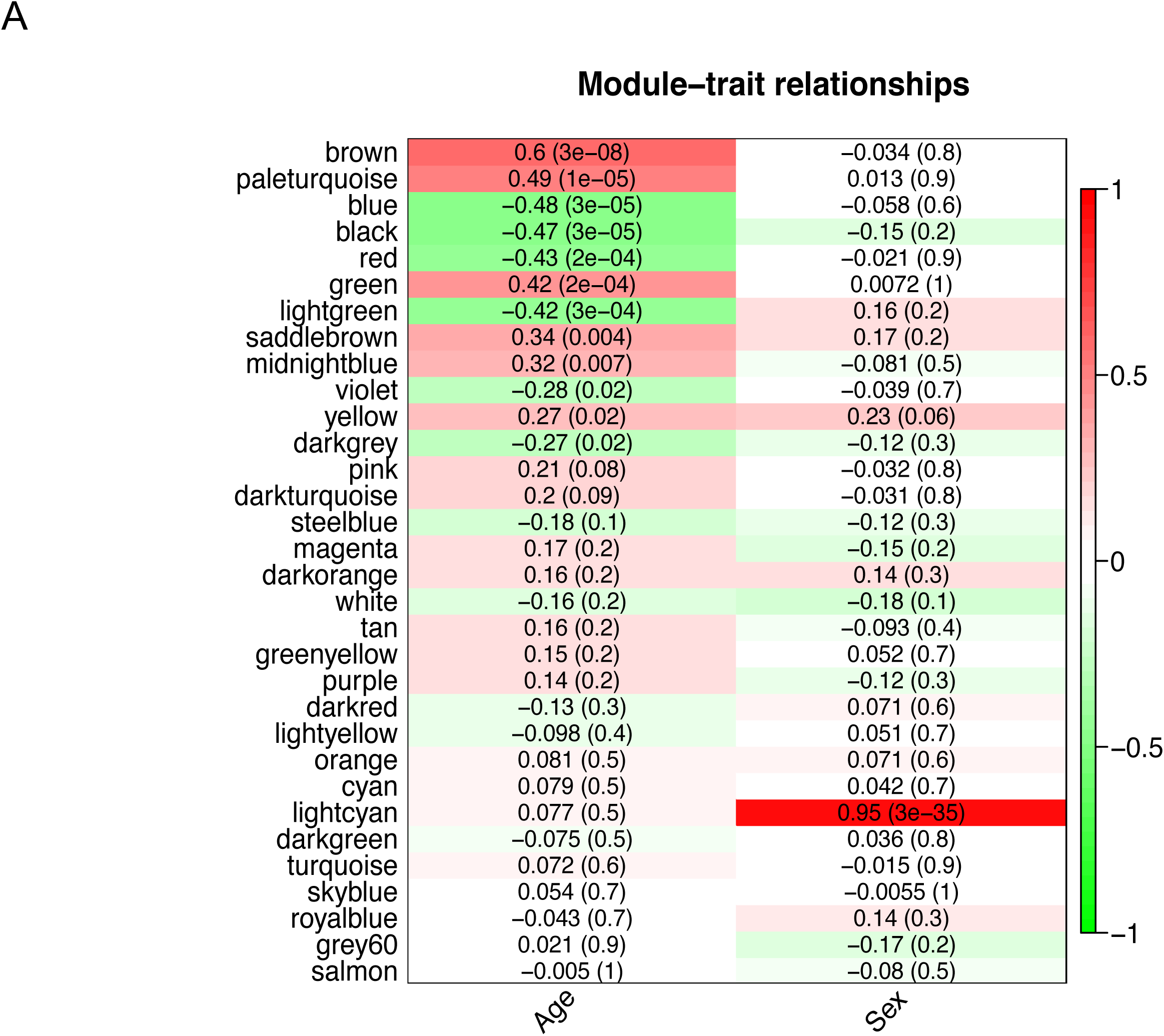
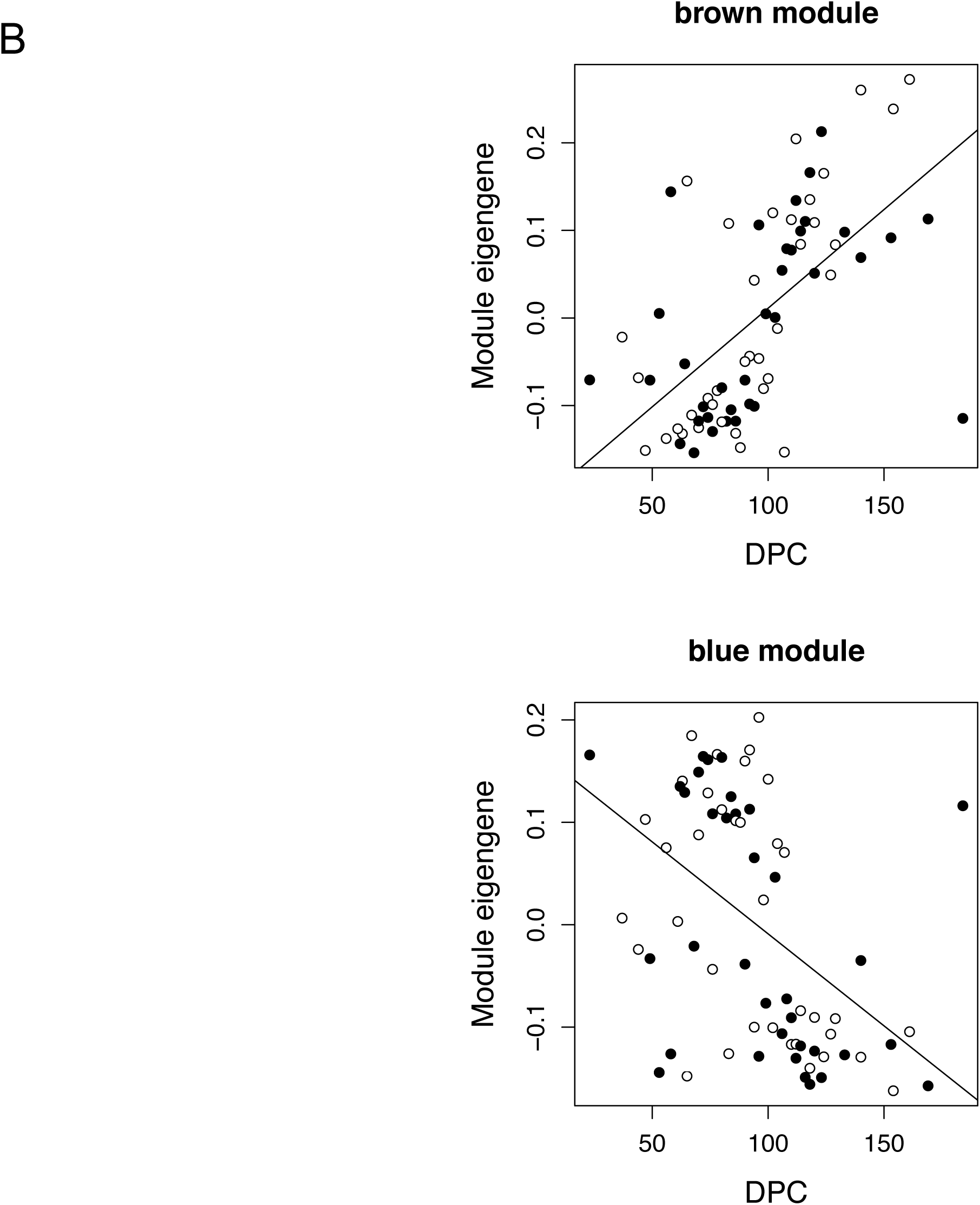
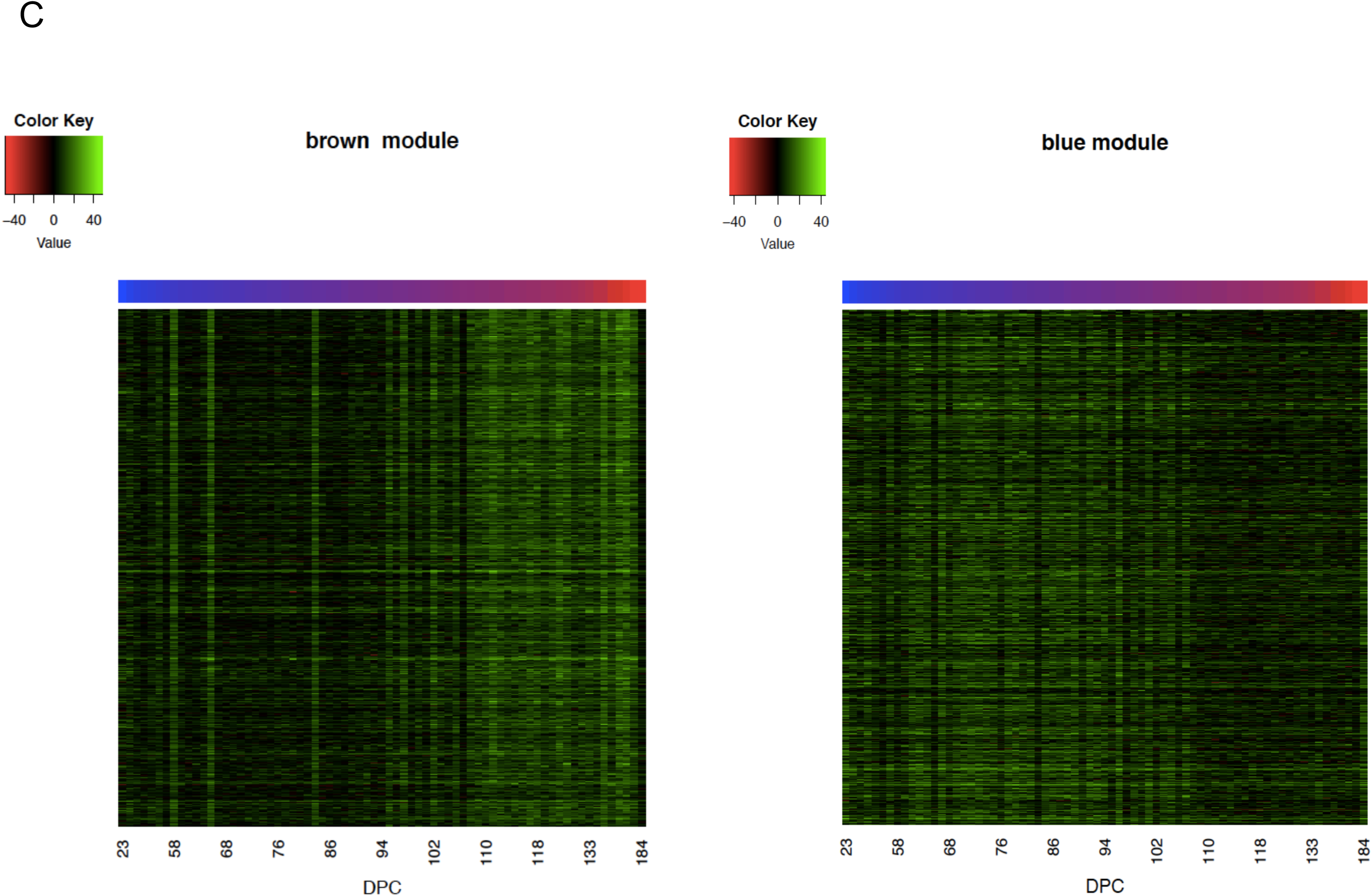
Modules of co-hydroxymethylated loci in the developing human brain. (A) Heat-map representing the correlation between module eigenvalues (ME) and the samples traits of fetal age (DPC) and sex. Each row represents a module, as indicated on the y-axis, and each column a trait. As shown in the color scale bar, strong positive correlation is indicated by dark red, strong negative correlation is indicated by dark green, and white indicates no correlation. Each cell contains the corresponding correlation and *P*-value given in parentheses (see also **Supplementary Table 15**). (B) Shown is the association between the ME and DPC for the top-ranked co-hydroxymethylated modules associated with brain development. The ‘brown’ module is the most positively associated with human fetal brain development (corr = 0.60, *P* = 3E- 08). The ‘blue’ module is the most negatively associated with human brain development (corr = -0.48, *P* = 3E-05). (C) Heat-map of the top 1,000 probes ranked by module membership in the ‘brown’ and ‘blue’ modules, showing coordinated changes in 5hmC across brain development. Color corresponds to the level of 5hmC at each probe.

### DNA hydroxymethylation quantitative trait loci

Building on our recent work identifying genetic effects on 5mC in the fetal brain[36], we explored genetic influences on levels of 5hmC by performing a genome-wide scan for 5hmC quantitative trait loci (hmQTL). Although we were not highly-powered to detect hmQTLs given the relatively small sample size and relative paucity of 5hmC, we identified 23 hmQTLs at a Bonferroni significance threshold (*P*-value < 2.2E-13) associated with 5hmC with a median effect size of 6.99% per allele (**Supplementary Table 17**, Fig. 5), and 77 hmQTLs at a more relaxed discovery threshold of *P*-value < 1E-11 (**Supplementary Table 18**). In contrast, we identified 13,480 mQTLs at a Bonferroni significance threshold (*P*-value < 1.6E-13) for 493 distinct sites, and 24,286 mQTLs at a discovery threshold of *P*-value < 1E-11 at 860 distinct sites, in the same samples, reflecting the higher abundance of methylated sites in the genome. Three sites were part of 11 Bonferroni significant QTLs for both modifications, all showing the same direction of effect. Of the 13,480 mQTL identified 1,383 (10.3%) were associated with probes that did not show any detectable levels of 5hmC, and 7,652 (56.8%) showed no evidence of a genetic effect on 5hmC (*P*-value > 0.05). 11,910 (98.5%) mQTLs were associated with significant (*P*-value < 0.05/12,097) heterogeneous genetic effects across 5mC and 5hmC.

**Figure 5:**
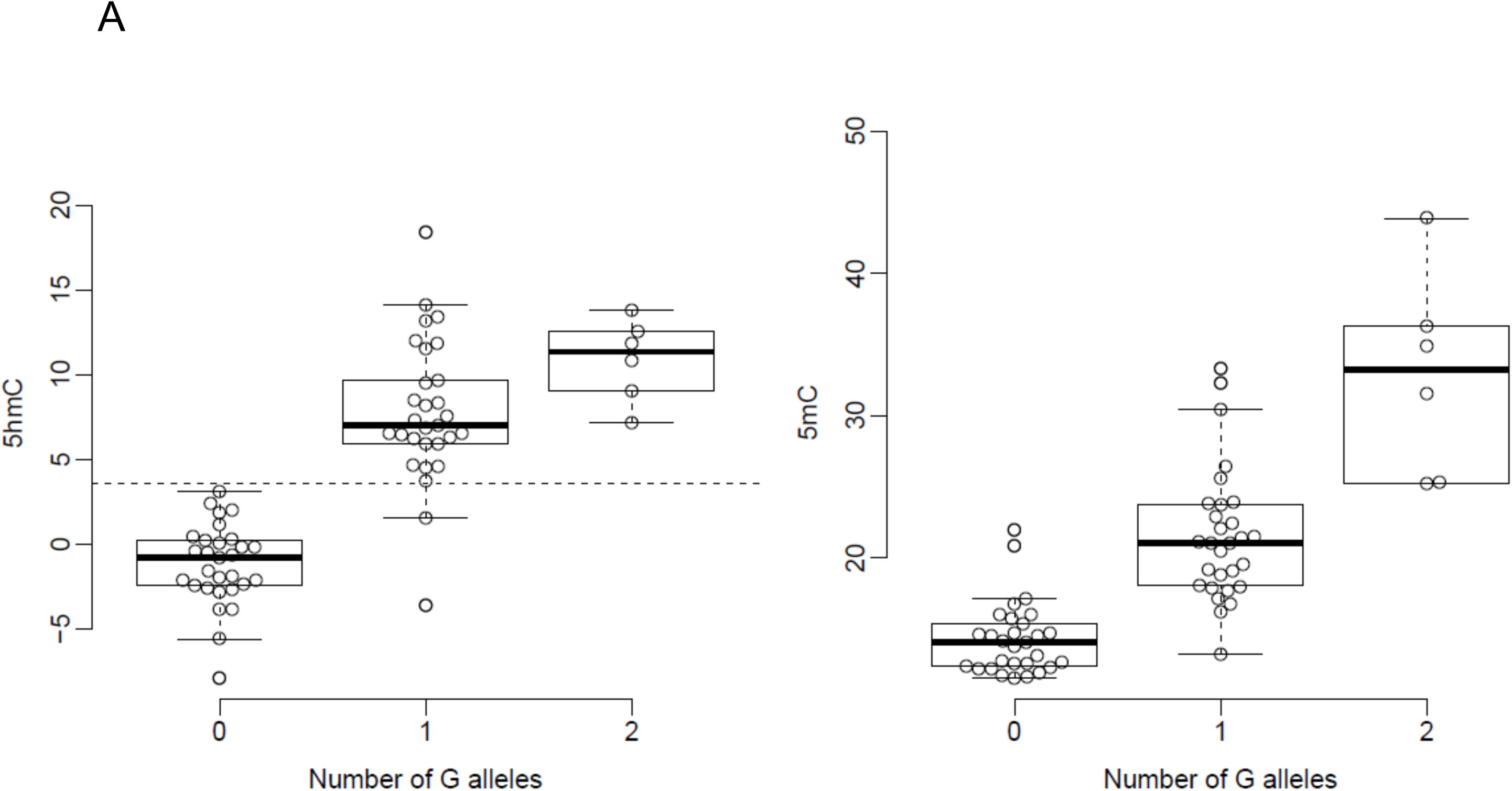
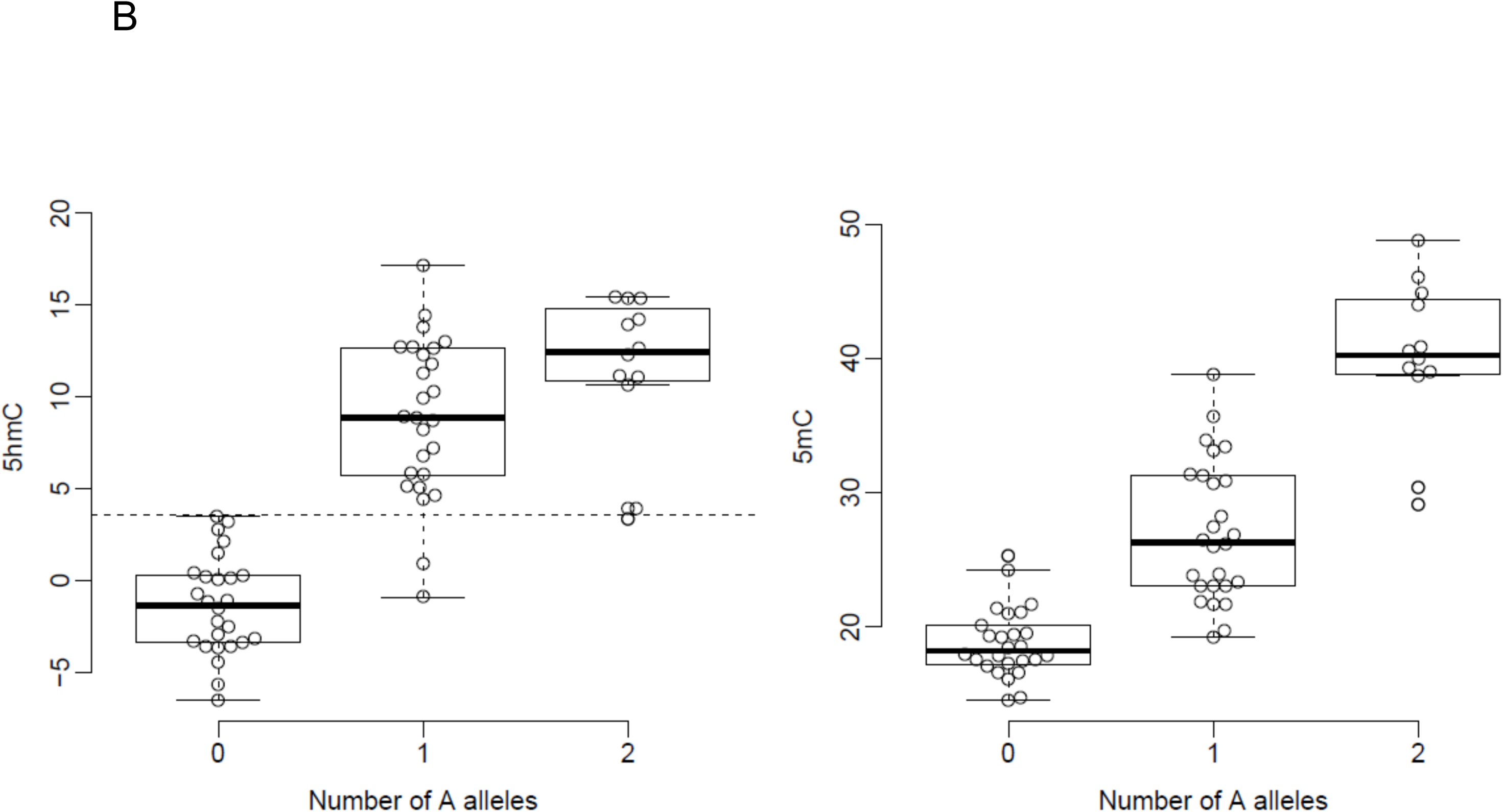
Examples where 5hmC at specific sites is influenced by genetic variation. (A) Genetic variation at rs11018924 influences both 5hmC (*P* = 4.67E-15) and 5mC (*P* = 9.73E-15) at cg26138821 in the fetal brain, with both modifications being positively associated with the minor allele (G). (B) Genetic variation at rs2788655 influences both 5hmC (*P* = 6.88E-12) and 5mC (*P* = 1.20E-10) at cg10523679, with both modifications positively associated with the minor allele (A).

## DISCUSSION

In this study, we examined neurodevelopmental changes in 5hmC in 71 human fetal brain samples spanning 23 to 184 DPC. To our knowledge, this represents the most extensive study of 5hmC in the human brain and is the first to examine changes in 5hmC across neurodevelopment. We identify highly significant changes in 5hmC during human brain development at multiple sites, and modules of co-hydroxymethylated loci associated with fetal age that are significantly enriched in the vicinity of genes involved in neurodevelopmental processes. We highlight a complex relationship between patterns of 5hmC and 5mC across brain development; while some loci are characterized by parallel changes in both modifications, other sites are characterized by opposite patterns. Strikingly, in contrast to 5mC, levels of 5hmC on the X-chromosome are lower in females than males. Finally, although hmQTLs are considerably less widespread than mQTLs in the fetal brain, we highlight examples where 5hmC is associated with genetic variation.

Consistent with previous observations in young neurons[18, 19, 27], we observe overall low levels of global 5hmC in the human fetal brain. Examining the genomic distribution of 5hmC highlighted an inverse correlation between mean 5hmC and CpG density, with a depletion of 5hmC in promoter regions and enrichment in gene bodies, relative to global 5hmC levels, concordant with previous observations in adult human brain[2, 19, 28-30, 37]. The chromosomal distribution of 5hmC was relatively consistent, although a depletion of 5hmC was found on the X-chromosome relative to autosomes in both males and females, and this deficit was particularly striking in females. These data concur with previous studies reporting a deficit in X-chromosome 5hmC[2, 19, 28], suggesting a possible role in dosage compensation for X-linked genes. Interestingly, there was reduced X-chromosome 5hmC in females compared to males, in stark contrast to 5mC which is associated with silencing of the inactive allele. 5hmC is hypothesized to be associated with transcriptionally permissive chromatin states; whilst 5mC is often associated with gene repression, 5hmC facilitates transcription through contributing to open chromatin[38, 39]. During synaptogenesis, for example, while 5mC accumulates in heterochromatin, 5hmC becomes enriched in euchromatin, consistent with its co-localization with PolII which mediates RNA transcription[20].

It has not yet been fully established whether 5hmC is a stable epigenetic mark that contributes to the regulation of genomic function, or whether it is a transient intermediate marking sites of active demethylation. Our analysis of the relationship between 5mC and 5hmC across brain development in the same samples revealed a complex relationship at individual probe sites. At a number of loci, levels of 5hmC increased or decreased in parallel with 5mC, although other locations were characterized by changes in the opposite direction during brain development. This stable accumulation or depletion of 5hmC at specific sites suggests it is not merely a transient intermediate of active DNA demethylation, providing further evidence to support a functional role for this modification in brain development.

There are several limitations to this study that should be considered. First, although this represents the first systematic analysis of 5hmC across multiple stages of human fetal brain development, legal restrictions on later-term abortions precluded the assaying of brain samples from later stages. Second, because the brain tissue was acquired frozen it was processed as a composite of brain regions and cell types and our data therefore reflect predominant changes in the developing human brain as a whole. Third, although the Illumina 450K platform provides accurate quantification at single-base resolution with probes associated with 99% of genes and 96% of CGIs, the array targets < 2% of the CpG sites in the human genome, and probes are not equally distributed across all genomic features. To calculate 5hmC it is necessary to run two arrays per sample (following BS- and oxBS-conversion), potentially increasing technical variability. However, we found a very strong concordance between the BS-converted DNA data and our previous study of total DNA modifications in the fetal brain. This study also highlights the potential for 5hmC to confound DNA modification data produced using BS-converted DNA, the standard method for profiling 5mC. As costs reduce, future studies can apply sequencing-based genomic profiling technologies to more thoroughly interrogate DNA modifications across the genome. To provide insight regarding the cause and functional consequences of epigenetic developmental dynamics, future studies should take a multi-omic approach and integrate genomic, transcriptomic and proteomic data sets; in particular, it will be important to integrate data about other epigenetic marks, including alternative DNA modifications (i.e. 5-fluorocytosine and 5-carboxylcytosine) and histone modifications with our 5mC and 5hmC data.

## CONCLUSIONS

We identify dynamic changes in 5hmC occurring throughout the genome during human fetal brain development. There are discrete modules of co-hydroxymethylated loci associated with brain development that are significantly enriched for genes involved in neurodevelopmental processes. There is a complex relationship between levels of 5mC and 5hmC across brain development, with notable sex differences on the X-chromosome. Finally, we show that 5hmC at specific loci is associated with genetic variation. Together, these data further highlight the role of dynamic epigenetic processes during brain development, with potential implications for understanding the origins of neurodevelopmental health and disease.

## METHODS

### Methodological overview

DNA modifications (DNA methylation (5mC) and DNA hydroxymethylation (5hmC)) were assessed in 72 human fetal brain samples (36 male; 36 female), spanning 23 to 184 days post-conception (DPC). Genomic DNA was isolated from fetal brain tissue acquired frozen from the Human Developmental Biology Resource (HDBR) (http://www.hdbr.org) and MRC Brain Banks Network (http://www.mrc.ac.uk/research/facilities/brain-banks/access-for-research) under strict ethical regulations, as previously described[3]. DNA modifications and DNA methylation were quantified with the Illumina Infinium HumanMethylation450 BeadChip, as previously described[26, 40]. Two pre-treatments, BS and oxBS conversion, and two Illumina 450K arrays were run for each sample. Following quality control procedures, preprocessing and normalisation, as previously described[40], DNA modification and DNA methylation beta values were generated, and a DNA hydroxymethylation value calculated from these measures. An overview of the method for the quantification of DNA hydroxymethylation is provided in **Supplementary Fig. 14**.

### Human fetal brain samples

Human fetal brain tissue was acquired from the Human Developmental Biology Resource (HDBR) (http://www.hdbr.org) and the MRC Brain Banks network (http://www.mrc.ac.uk/research/facilities/brain-banks/access-for-research). Ethical approval for the HDBR was granted by the Royal Free Hospital research ethics committee under reference 08/H0712/34 and Human Tissue Authority (HTA) material storage license 12220; ethical approval for the MRC Brain Bank was granted under reference 08/MRE09/38. The developmental age of each sample was determined using Carnegie Staging for embryonic (defined as ≤ 56 DPC) samples and foot and knee to heel length measurements for fetal (defined as ≥ 57 DPC) samples. No additional phenotypic or demographic information was available for any of the samples used in this study. An overview of the samples is given in **Supplementary Fig. 15** and **Supplementary Table 19**. Genomic DNA was isolated using standard phenol-chloroform procedures and quantified using the Qubit^®^ BR dsDNA assay with the Qubit^®^ Fluorometer (Life Technologies, Thermo Fisher Scientific). Sample sex was determined via PCR amplification of a region near the amelogenin gene (*AMELY*) that produces different sized PCR products for the X and Y Chromosomes, (977 and 788 bp, respectively)[41]. Following DNA modification profiling, sex was confirmed using multidimensional scaling plots of microarray probes on the X Chromosome.

### Quantification of DNA methylation and DNA hydroxymethylation using the Illumina 450K array

Samples were randomized with respect to DPC and sex to avoid batch effects throughout all experimental procedures. For each sample two 500 ng aliquots of genomic DNA were subject to parallel oxBS and BS conversion (**Supplementary Fig. 14**) using the TrueMethyl^®^ 24 kit (CEGX, Cambridge, UK). Sample recovery was assessed using the Qubit^®^ ssDNA assay, with a final sample concentration of ≥ 20 ng/μl considered optimal for further analysis. A digestion control assay was used to qualitatively confirm successful oxBS and BS conversion (**Supplementary Fig. 16**). Total levels of DNA modification (5mC + 5hmC, BS-converted DNA) and DNA methylation (5mC, oxBS-converted DNA) were quantified using the Illumina HumanMethylation450 BeadChip (Illumina) scanned on an Illumina HiScan (Illumina). BS- and oxBS-converted DNA samples from each individual were processed together at all stages.

### Data pre-processing and quality control

GenomeStudio (Illumina) was used to extract signal intensities for each probe, generating a final report that was imported into the R statistical environment 3.0.2 (http://www.r-project.org)[42] using the methylumi package (http://www.bioconductor.org/packages/release/bioc/html/methylumi.html). Sample identity was confirmed using multidimensional scaling to confirm concordance between the expected and reported sample sex, and via correlation of the beta values for the 65 single nucleotide polymorphism (SNP) probes for each BS- and oxBS-converted sample pair. Data quality control and preprocessing were performed using the *wateRmelon* package as described previously[40]. Stringent filtering of the prenormalized Illumina 450K data was performed using the *pfilter* function; CpG sites with a beadcount of < 3 in 5% of samples were removed (n = 838) and CpG sites with a detection *P*-value > 0.05 in 1% of samples (n = 19,163), additionally one sample having 5% of sites with a detection *P*-value > 0.05 was removed. Finally, cross-reactive probes and polymorphic CpGs, as detailed in the Illumina annotation file and identified in recent publications[43, 44] were removed (**Supplementary Table 20**), leaving 411,325 probes for initial analysis (including n = 9275 X-chromosome and n = 15 Y-chromosome probes). Information for samples included in the final dataset (n = 35 males [age range = 23 to 184 DPC]; n = 36 females [age range = 37 to 161 DPC]) is shown in **Supplementary Table 19**. DNA modification (BS-converted) and DNA methylation (oxBS-converted) data were normalized separately using the *dasen* method within the *wateRmelon* package[40]. Plotting the normalised beta density for both the BS- and oxBS-converted samples revealed a bimodal distribution for both pre-treatment approaches (**Supplementary Fig. 17**). The BS-converted samples showed a clear shift to the right of the oxBS-converted samples, corresponding to combined quantification of both 5mC and 5hmC. Normalized beta values for the oxBS-converted samples (5mC) were subtracted from the normalized beta values for the corresponding BS-converted samples (5hm + 5hmC) to calculate a ‘Δ beta’ value corresponding to the level of DNA hydroxymethylation at each probe. As expected, the beta value density distribution was positively skewed (**Supplementary Fig. 18**). As previously described[26], a threshold for defining “detectable” 5hmC was calculated from the 95^th^ percentile of the negative beta values (which result from technical noise), and beta values > 0.0357 were considered to represent reliably detected levels of 5hmC. Probes not passing this threshold in at least one sample (n = 103,501 of the 411,325 sites passing quality control) were removed (**Supplementary Table 1**); leaving 307,824 probes (298,972 = autosomal) for further analysis.

### Identification of differentially modified position as (DMPs) and regions (DMRs) associated with human brain development

A multiple linear regression model was used to identify changes in 5mC and 5hmC at specific sites associated with brain development and gender. Autosomal probes (n = 298,972) were considered significantly associated with brain development if they passed a Bonferroni-corrected significance threshold of *P*-value < 1.67E-07. Autosomal and X-chromosome probes (n = 307,810) were considered to be significantly associated with sex if they passed a Bonferroni-corrected significance threshold of *P*-value < 1.62E-07. To identify DMRs, we used the Python module *comb-p*[32] to group spatially correlated DMPs (seed *P*-value < 1.00E-04, minimum of 3 probes, Šidák-corrected *P*-value < 0.05) at a maximum distance of 500 bp. DMR *P*-values were corrected for multiple testing using Šidák correction which corrects the combined *P*-value for n_a_/n_r_ tests, where n_a_ is the total number of probes tested in the initial dataset and n_r_ the number of probes in the given region.

### Weighted gene co-modification network analysis

Weighted gene co-modification network analysis (WGCNA) was used to identify modules of highly co-hydroxymethylated probes[35]. Modules were identified from their pairwise correlations, using a signed block-wise network construction at power 5 with a merging threshold of 0.30 and n = 1000 minimum probes. Modules were assigned a color name, and a weighted average-hydroxymethylation profile, known as a module eigengene (ME), was calculated for each. Correlations between the MEs and the phenotypic traits (DPC, sex) were used to identify modules associated with these traits. The relationship between each probe and each module was assessed through calculation of module membership (MM), the absolute correlation between a probe’s DNA hydroxymethylation status and the ME, allowing the identification of the subset of probes with the strongest membership to each module. Probe significance (PS), the absolute value of correlation between each probe and phenotypic trait, was calculated to quantify the association of probes with DPC. To assess the biological meaning of modules associated with DPC and sex, genes associated with the top 1000 probes ranked by MM were extracted and assessed using pathway and gene ontology analysis.

### Gene ontology analysis

A gene list was derived from the DMPs using Illumina's gene annotation. This annotation, which comes via UCSC, is based on overlap with RefSeq genes plus 1500 bp of upstream sequence. Where probes were not annotated to any gene (i.e. in the case of intergenic locations) they were omitted from this analysis, and where probes were annotated to multiple genes all were included. A logistic regression approach was used to test if genes in this list predicted pathway membership, while controlling for the number of probes that passed quality control (i.e. were tested) annotated to each gene. Pathways were downloaded from the GO website (http://geneontology.org/) and mapped to genes including all parent ontology terms. All genes with at least one 450K probe annotated and mapped to at least one GO pathway were considered. Pathways were filtered to those containing between 10 and 2000 genes. After applying this method to all pathways, the list of significant pathways (*P*-value < 0.05) was refined by grouping to control for the effect of overlapping genes. This was achieved by taking the most significant pathway, and retesting all remaining significant pathways while controlling additionally for the best term. If the test genes no longer predicted the pathway, the term was said to be explained by the more significant pathway, and hence these pathways were grouped together. This algorithm was repeated, taking the next most significant term, until all pathways were considered as the most significant or found to be explained by a more significant term.

### DNA methylation and DNA hydroxymethylation quantitative trait loci

Genotype data was available for 64 of the 71 samples; these were assessed for DNA methylation quantitative trait loci (mQTL) and DNA hydroxymethylation quantitative trait loci (hmQTL). The genetic data was profiled on the Illumina HumanOmniExpress BeadChip (Illumina) chip and imputed as described previously[36]. SNPs were then filtered with PLINK[45] excluding variants with > 1% missing values, Hardy-Weinburg equilibrium *P*-value < 0.0001 or a minor allele frequency of < 5%. Subsequently, SNPs were also filtered so that each of the three genotype groups with 0, 1, or 2 minor alleles (or two genotype groups in the case of rarer SNPs with 0 or 1 minor allele) had a minimum of 5 observations. An additive linear model was fitted using MatrixEQTL[46] to test if the number of alleles (coded 0, 1, or 2) predicted 5mC or 5hmC at each site, including covariates for age, sex, and the first two principal components from the genotype data. To test for a significantly different genetic effect on 5hmC compared to 5mC, all significant mQTL and hmQTL were subsequently tested for heterogeneous effects. To this end, a multi-level model was fitted across the data for both modifications using the R packages lme4[47] and lmerTest[48]. In these models genotype, age, sex, the first two principal components and an indicator variable to distinguish 5mC from 5hmC measurements were included as fixed effects, and individual was included as a random effect. Two models with and without an interaction term between genotype and the modification indicator variable were fitted for each QTL and the heterogeneity *P*-value was calculated from an ANOVA comparing these two models.

## DECLARATIONS

### ETHICAL APPROVAL

Ethical approval for the HDBR was granted by the Royal Free Hospital research ethics committee under reference 08/H0712/34 and Human Tissue Authority (HTA) material storage license 12220; ethical approval for the MRC Brain Bank was granted under reference 08/MRE09/38.

### AVAILABILITY OF DATA

Raw and normalized Illumina 450K methylation data has been submitted to the NCBI Gene Expression Omnibus (GEO; http://www.ncbi.nlm.nih.gov/geo/) under accession number GSE94014. A searchable database of our fetal brain epigenomics data is available at http://epigeneticsessex.ac.uk/fetalbrain2/.

## ACKNOWLEDGEMENTS

This project was supported by a Brain & Behavior Research Foundation Distinguished Investigator Award to J.M. H.S. was supported by an MRC PhD studentship. The human embryonic and fetal material was provided by the Joint MRC (grant #G0700089)/Wellcome Trust (grant #GR082557) Human Developmental Biology Resource (http://www.hdbr.org).

## AUTHOR CONTRIBUTIONS

J.M. and N.B. conceived and supervised the study. J.M. obtained funding. H.S. performed the laboratory work, data analysis and bioinformatics. E.H. provided analytical support and performed analysis. L.S. provided analytical support and developed the website database resource. H.S., E.H. and J.M. drafted the manuscript. All of the authors read and approved the final submission.

## COMPETING INTERESTS

The authors declare no competing financial interests.

## FIGURE LEGENDS

**Figure 1:5hmC is highly dynamic across human fetal brain development.**

(A) The presence or absence of detectable 5hmC at individual sites is highly variable amongst the fetal brain samples profiled in this study. Shown is the proportion of probes with detectable 5hmC as a function of the fetal brain samples profiled. Only a small number of probes are characterized by ubiquitously detectable 5hmC in all samples, highlighting considerable inter-individual heterogeneity at individual probes. (B) The top-ranked dDHP characterized by increasing 5hmC with days post-conception (DPC) is cg25069807, annotated to *TK1* on chromosome 17 (P = 2.52E-09). (C) The top-ranked dDHP characterized by decreasing 5hmC with brain development is cg22960621, annotated to *NLGN1* on chromosome 3 (*P* = 1.42E-09). Males are depicted in black. The dashed line indicates values above which 5hmC is considered to be reliably detected (Δβ_BS-oxBS_ > 0.036). The full list of 2,181 dDHPs (*P* < 5.00E-05) is presented in **Supplementary Table 5.** (D) The top-ranked dDHR associated with fetal brain development spans a 685 bp region (Chr22: 46449430 – 46450115) encompassing the first exon of *C22orf26* and *LOC150381*. The dDHR is denoted by dashed vertical lines. Chromosomal coordinates correspond to human genome build GRCh37/hg19. See **Supplementary Table 8** for a full list of dDHRs associated with human brain development.

**Figure 2:There is a complex interaction between 5hmC and 5mC across human brain development.**

(A) Scatterplot showing the regression coefficients for 5hmC and 5mC for each of the 2,181 dDHPs (*P* < 5.00E-05). Nearly half of the dDHPs (n = 1059, 48.56%) are characterized by a significant interaction (*P* < 2.29E-05) between 5hmC and 5mC across brain development (significant sites are shown in red). Examples of sites at which fetal brain development is associated with (B) increases in both 5hmC and 5mC, (C) decreases in both 5hmC and 5mC, (D) an increase in 5mC but a decrease in 5hmC, (E) a decrease in 5mC but an increase in 5hmC and (F) a change in 5hmC but no alteration in 5mC. 5mC data shown as diamonds, 5hmC data shown as circles. Male samples are indicated by a filled diamond or circle.

**Figure 3:Levels of 5hmC in the human fetal brain are significantly lower on the X-chromosome in females than males.**

(A) Manhattan plot showing the distribution of sites characterized by different levels of 5hmC between males and females. *P*-value corresponds to association with sex. Points above the X-axis indicate higher 5hmC in females, with those below the X-axis indicating higher 5hmC in males. The red line represents a Bonferroni-corrected significance threshold (*P* < 1.62E-07) for the 307,810 autosomal and X chromosome probes analyzed. The blue line corresponds to our “discovery” *P*-value threshold (*P* < 5E-05). (B) The top-ranked sex-associated DHP characterized by greater 5hmC in males is cg07806797, annotated to ZNF185 on the X-chromosome (*P* = 7.19E-20). (C) The top-ranked sDHP characterized by greater 5hmC in females is cg2678647, annotated to USP11 on the X-chromosome (*P* = 3.64E-16). (D) Mean 5hmC across canonical gene features is shown for males and females (pink and blue, respectively) for autosomes (top panel) and the X-chromosome (bottom panel).

**Figure 4:Modules of co-hydroxymethylated loci in the developing human brain.**

(A) Heat-map representing the correlation between module eigenvalues (ME) and the samples traits of fetal age (DPC) and sex. Each row represents a module, as indicated on the y-axis, and each column a trait. As shown in the color scale bar, strong positive correlation is indicated by dark red, strong negative correlation is indicated by dark green, and white indicates no correlation. Each cell contains the corresponding correlation and *P***-**value given in parentheses (see also **Supplementary Table 15**). (B) Shown is the association between the ME and DPC for the top-ranked co-hydroxymethylated modules associated with brain development. The ‘brown’ module is the most positively associated with human fetal brain development (corr = 0.60, *P* = 3E-08). The ‘blue’ module is the most negatively associated with human brain development (corr = -0.48, *P* = 3E-05). (C) Heat-map of the top 1,000 probes ranked by module membership in the ‘brown’ and ‘blue’ modules, showing coordinated changes in 5hmC across brain development. Color corresponds to the level of 5hmC at each probe.

**Figure 5:Examples where 5hmC at specific sites is influenced by genetic variation.**

(A)Genetic variation at rs11018924 influences both 5hmC (*P* = 4.67E-15) and 5mC (*P* = 9.73E- 15) at cg26138821 in the fetal brain, with both modifications being positively associated with the minor allele (G). (B) Genetic variation at rs2788655 influences both 5hmC (*P* = 6.88E-12) and 5mC (*P* = 1.20E-10) at cg10523679, with both modifications positively associated with the minor allele (A).

